# Artificial Neurovascular Network (ANVN) to Study the Accuracy Vs. Efficiency trade-off in an Energy Dependent Neural Network

**DOI:** 10.1101/2021.02.16.431351

**Authors:** Bhadra S Kumar, Nagavarshini Mayakkannan, N Sowmya Manojna, V. Srinivasa Chakravarthy

## Abstract

**Abstract:** Artificial feedforward neural networks perform a wide variety of classification and function approximation tasks with high accuracy. Unlike their artificial counterparts, biological neural networks require a supply of adequate energy delivered to single neurons by a network of cerebral microvessels. Since energy is a limited resource, a natural question is whether the cerebrovascular network is capable of ensuring maximum performance of the neural network while consuming minimum energy? Should the cerebrovascular network also be trained, along with the neural network, to achieve such an optimum?

In order to answer the above questions in a simplified modeling setting, we constructed an Artificial Neurovascular Network (ANVN) comprising a multilayered perceptron (MLP) connected to a vascular tree structure. The root node of the vascular tree structure is connected to an energy source, and the terminal nodes of the vascular tree supply energy to the hidden neurons of the MLP. The energy delivered by the terminal vascular nodes to the hidden neurons determines the biases of the hidden neurons. The “weights” on the branches of the vascular tree depict the energy distribution from the parent node to the child nodes. The vascular weights are updated by a kind of “backpropagation” of the energy demand error generated by the hidden neurons.

We observed that higher performance was achieved at lower energy levels when the vascular network was also trained along with the neural network. This indicates that the vascular network needs to be trained to ensure efficient neural performance. We observed that below a certain network size, the energetic dynamics of the network in the *per capita energy consumption* vs. *classification accuracy* space approaches a fixed-point attractor for various initial conditions. Once the number of hidden neurons increases beyond a threshold, the fixed point appears to vanish, giving place to a line of attractors. The model also showed that when there is a limited resource, the energy consumption of neurons is strongly correlated to their individual contribution to the network’s performance.

**Author summary:** The limited availability of resources contributed to a significant role in shaping evolution. The brain is also no different. It is known to have tremendous computing power at a significantly lower cost than artificial computing systems. The artificial neural networks aim typically at minimizing output error and maximizing accuracy. A biological network like the brain has an added constraint of energy availability, which might force it to choose an optimal solution that provides the best possible accuracy while consuming minimum energy. The intricate vascular network which ensures adequate energy to the brain might be a systematically trained layout rather than a hard-wired anatomical structure. Through this work, we intend to explore how the artificial neural network would behave if it were made dependent on an energy supply network and how the training of the energy supply network would influence the performance of the neural network. Our model concluded that training of a vascular energy network is highly desirable, and when the size of the neural network is small, the energy consumed by each neuron is a direct readout on its contribution to the network performance.

## Introduction

Energy constraints are believed to play a vital role in shaping the evolution of the brain ^1,2^. Although the brain performs amazingly well in analog signal processing, it is interesting to note that it is not evolved to give maximum information transfer efficiency; instead, it compromises information transfer efficiency in a trade-off with energy efficiency^3^.

Although a majority of computational neuroscience models focus on modeling neural signaling, in recent years, there is a growing body of literature that addresses issues related to energy dynamics that underlies neural signaling, conceptualized as a separate field known as *neuroenergetics* ^4^. Neural signaling, which is associated with fluctuations in the membrane voltage, is supported by the current flowing through the ion channels. However, alterations in concentrations of the ionic species (Na^+^, K^+^, Ca^2+^) on either side of the neural membrane are corrected by the action of various pumps that utilize energy in the form of Adenosine TriPhosphate (ATP) (Chapters 5,6,7, Kandel et al., 2000). This ATP is replenished by oxidative phosphorylation and glycolysis using the oxygen and glucose received from the blood vessels. Proximal blood vessels dilate as a consequence of neural activity, which ensures adequate blood flow to fuel the neural activity ^7–9^.

The interaction between neurons and cerebral vessels is called neurovascular coupling. Many molecular pathways facilitate effective coupling between neurons and vessels. One important mechanism is the direct release of nitric oxide by the neurons ^10,11^. Nitric oxide is a vasoactive substance that diffuses to nearby vessels, causing their dilation. Even though the forward influence from the neuron to a vessel is widely studied, particularly in the context of understanding functional neuroimaging ^12^, the retrograde influence from the vessels to the neurons, responsible for converting oxygen and glucose to ATP is still not completely understood. One widely accepted theory is that the glucose transported to the neurons are picked up directly by neurons and used as a substrate to carry out oxidative phosphorylation, which is the major contributor of ATP, the energy currency used to fuel the action of ionic pumps ^13–15^. Another exciting study still debated ^16–20^ is the astrocyte-neuron lactate shuttle theory ^21^, which posits an important role to astrocyte in mediating neurovascular coupling. Astrocytes convert the glucose released from the cerebral vessels to lactate and provide it to the neurons, where it is converted into pyruvate and ultimately into ATP.

Though there have been efforts to model the elaborate bidirectional signaling underlying neural energetics at the single neuron level ^22–26^, there is also an obvious need to study energetics at a network level. There is a growing awareness that impaired neuroenergetics is involved in several important brain disorders ^27–31^. Abnormally high energy consumption levels have been linked to the idiopathic loss of cells in Substantia nigra in Parkinson’s disease ^32–34^. Mitochondrial dysfunction in the neurons of CA1 and CA3 in the hippocampus is linked to cellular pathology underlying Alzheimer’s disease ^35,36^. There have been bold proposals that metabolic impairments are the common underlying cause behind all forms of neurodegeneration ^37^. Disruptions in neurovascular coupling evidently accompany the brain disorders like stroke and vascular dementia ^38–42^. It would be beneficial to develop computational models that provide insights into the genesis of pathological neuroenergetics in the aforementioned diseases. But if the models of neuroenergetics at the single-unit level are so complicated, it would not be a pragmatic enterprise to extend them as they are to a network level and study neuroenergetics in an extremely detailed fashion at a network level.

Therefore, to study neuroenergetics at the network level, the description at the single-unit level must be appropriately simplified. Description of neural energetics in terms of a large number of molecular metabolic substrates (e.g., ATP, pyruvate, lactate, glucose) makes the model conceptually opaque. The study of energetics in engineering is greatly facilitated by the unified view of energy that has been worked out in physics over the centuries. Behind the innumerable forms of energy found in nature (mechanical, electrical, chemical, thermal, etc.), energy is one, denoted by common units (Joules). However, such an elegant, intellectually satisfying, and unified view of energy transformations in neuroenergetics, though desirable, is a far-off goal considering the current state of neuronal modeling.

The development of simplified neural network models in the ’80s had led to the connectionist revolution. Although more complex modeling approaches were available at that time – like the Hodgkin-Huxley model, the cable equation, and dendritic processing - networks constructed using simple sigmoidal neurons have succeeded in providing insights into a wide variety of phenomena in psychology and cognitive science in addition to artificial engineering domain ^43,44^.

Likewise, one may envisage that the development of simplified neuroenergetic models at the network level may give valuable insights into possible optimal energy utilization strategies of the brain. There have been efforts to construct simplified neuro-energetic models in recent times at the single neuron level. Some of these efforts assume that the effect of energy supply to a neuron can be expressed in abstract terms as regulating the neuron’s threshold of activation: higher (lower) energy leads to a smaller (higher) threshold ^45,46^.

By adopting such a simplified depiction of the dependence of neural function on energy, we construct a novel class of neural networks known as Artificial Neuro-Vascular Networks (ANVNs). In these networks, a separate vascular tree caters to the energy requirements of a neural network. Error gradient, which is usually used to update the weights of the neural network, is propagated, from the neural network, up the vascular tree to update the “strengths” of various branches of the vascular tree. The efficiency of energy utilization of the network, evaluated in terms of energy consumed to achieve a given level of output performance, is studied.

The outline of the paper is as follows. In the first half of the work, we bring out the importance of training the energy network of an energy-dependent network, in this case, the highly energy-dependent biological neural network. The energy network, in this case, is the vascular network that provides adequate nourishment to the brain tissues. We connect a neural network with a trainable vascular network to form an ANVN. The ANVN is then studied under three training regimes of the vascular tree (i) Leaving the vascular tree untrained, (ii) Training the vascular tree sequentially after first training the neural network, and (iii) Simultaneously training both neural and vascular networks. The improved performance in terms of accuracy and energy efficiency during simultaneous training (regime iii) establishes the need to train the vascular network. The energy provided to the network is variable, and the network takes up whatever energy is given to it.

In the second half of the work, we modify the ANVN to ANVN_R (ANVN with reservoir) such that it takes only the required energy and rejects the excess energy. This modification brings out the notion of an optimum network size where the energy efficiency can be maximized. The importance of the network size in maintaining its robustness to the initial availability of energy is also studied in the same section. We also look at how transfer learning would manifest in the vascular weight modification and observes that the weights representing the microvasculature undergo the maximum modification.

In the last section, we explore how bringing an explicit energy constraint in the cost function, as a form of regularization, would affect the network behavior. Finally, we study whether the energy consumed by a neuron reflects its contribution to the network performance. The model shows that such a correlation between energy consumption and the contribution of the neuron to the network exists only when the network size is small.

## Materials and methods

The Artificial Neuro Vascular Network (ANVN) is designed with the intention to explore the characteristics of an energy-dependent neural network. In ANVN, we establish a bidirectional coupling between a simple feedforward neural network like the Multi-Layer Perceptron (MLP) and a vascular network model. The bidirectionality, in this context, refers to the following flows: the forward flow of energy from the vascular network to the neural network and the feedback flow of energy demand error from the neural network to the vascular network. fig. (1) shows the schematic of the model. In fig.1, ANVN with reservoir (ANVN_R) is shown. For the initial part of the study, we consider ANVN, which has only the root node and no reservoir. The source and reservoir in fig. 1 are the part of the modified ANVN with reservoir (ANVN_R), which comes in the later part of the study. The hidden layer of the neural network is shown by the yellow box, and the yellow circles indicate the neurons. As shown in the figure, each neuron receives energy from the leaf nodes of the vascular network (green circles), and the energy available at each leaf node depends on the weights of the vascular tree indicated by *U_y,x_* connecting nodes x and y. During the forward pass in MLP, the bias of the hidden neurons is decided by the amount of energy available at the neuron. During backpropagation, the gradient in biases of hidden layer neurons is converted to gradients in energy and is propagated along the vascular tree from the leaf node to the root node (till source node for ANVN_R). The weights of the vascular tree are updated depending on this energy gradient.

**Figure.1.**
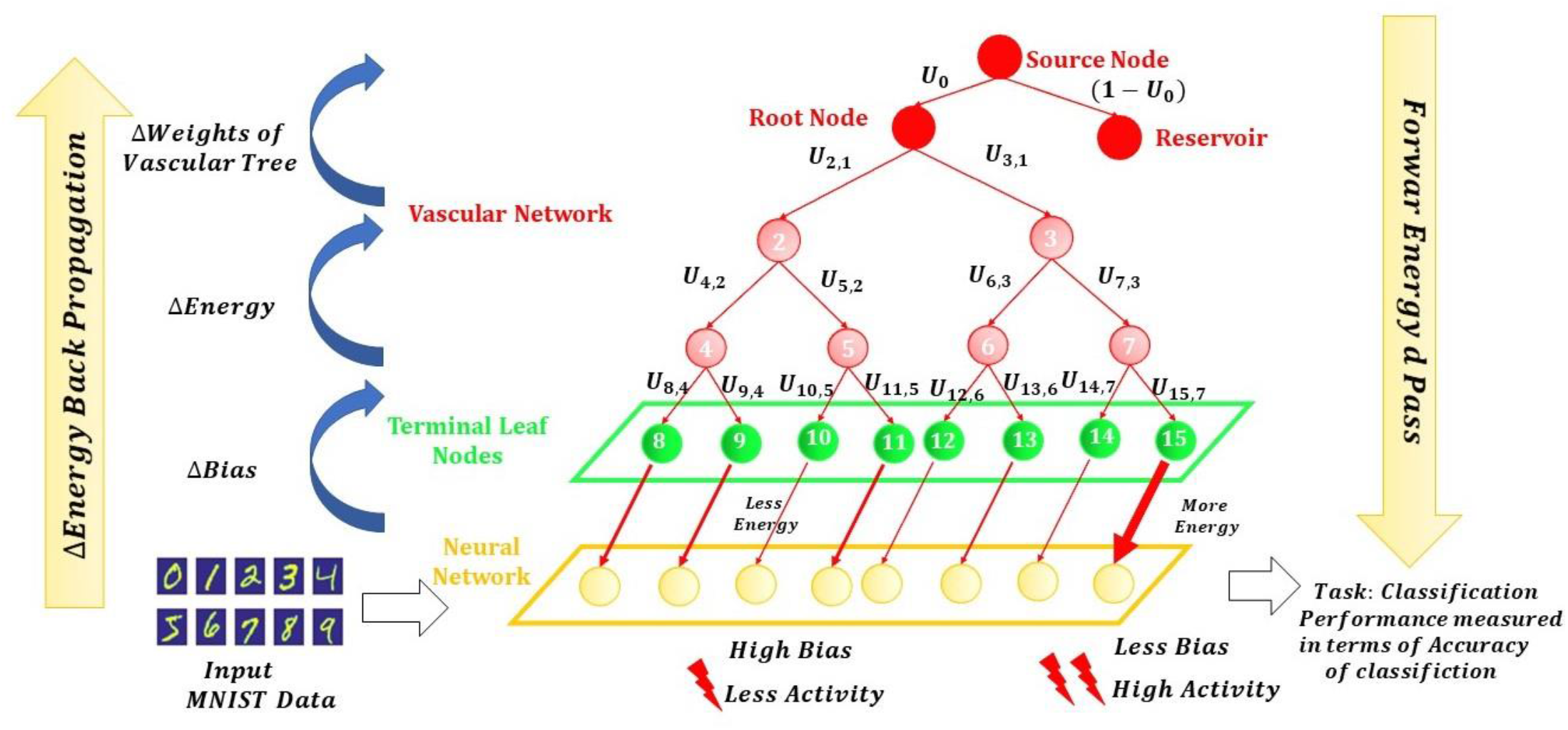
The schematic representation of ANVN_R

### The Vascular Network

The vascular network has a tree structure that begins at a root node (*R*) and branches uniformly depending on the predefined branching factor (≤ *k* branches at every junction) until it reaches the leaf nodes. The leaf nodes (F) supply energy to the hidden neurons of the MLP in a one-to-one fashion, and therefore equal in number to the number of hidden neurons (V). Hence the number of leaf nodes is initialized to be equal to the number of hidden neurons (F=N). The vascular tree is defined by fixing *k and N* apriori. The total number of levels in the tree (*L*) is calculated as,

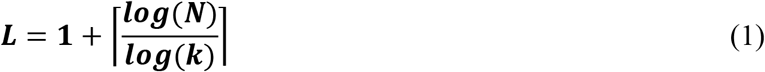

The total number of nodes *(T_n_*) in the tree is calculated as follows,

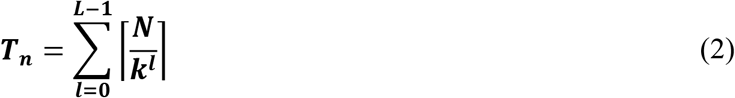

In case the number of neurons *N,* and consequently the number of leaf nodes (F) is not a power of branching factor (*k*), considering the level of the root node (R) as *l* = 1, each level gives rise to 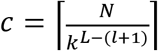 children in such a way that first 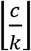 nodes have *k* branches, and the remaining nodes will have < *k* branches 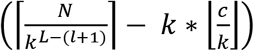. The tree structure is represented using the adjacency matrix (A), which is a two-dimensional matrix of size *T_n_* × *T_n_*. An entry *A_yx_* of the matrix is unity if there exists an edge from node *x* to node *y.*

The input energy source is connected to the root node. Energy from the root (*E_s_*) flows down the tree from the root node to the leaf nodes and from there to the hidden neurons. Each parent node (*x*) is connected to its child node (*y*) by a weighted connection 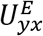. This weight defines the amount of energy transferred from the parent node to the child node. Since a given parent node can have up to *k* branches, each weight connection is absolute normalized with respect to its sibling branches (≤ *k*) to ensure the conservation of energy.

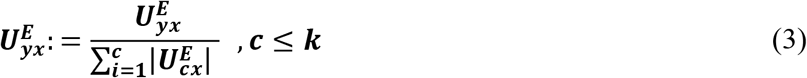

The vascular weights then represent the fraction of energy that flows through a branch compared to its sibling branches. The vascular weight matrix (*U^E^*) of size *T_n_* × *T_n_* is obtained by replacing the unit values of the adjacency matrix (A) with the weight of that edge. The energy of all parent nodes of level *l* is projected on the weight matrix to obtain the energy of the children nodes, which constitute the nodes of level *l*+1.The energy vector (***E_L_***) is of size *T_n_* × 1 and represents the energy distribution across all the nodes of the tree. ***E_L_*** has to be calculated recursively by updating ***E_x_*** at each level as shown in equation (4), starting from the level of the root node to the level of terminal leaf nodes in order to find the energy distribution at all the nodes. ***E_x_*** is initialized such that the energy at the root node is equal to the available energy *E_s_*. This initialized vector which is used as (***E***_1_) has nonzero value only at the first index, which represents the root node.

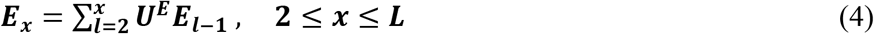

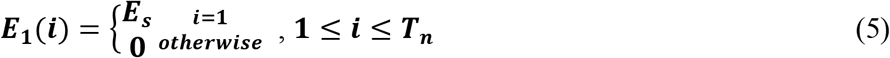

The vascular weights are initially randomly distributed, resulting in a random distribution of energies at leaf nodes. The energy available at each leaf node is used to calculate that hidden neuron’s bias, as described in equation 7 below.

### The Neural Network

The neural network used in this model is an MLP with a single hidden layer. For the first half of the study, the number of neurons in the hidden layer is fixed as N=512, and the performance of ANVN is studied as input energy is varied. A weighted connection 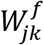 connects neuron ‘k’ of input layer to neuron ‘j’ of the hidden layer and 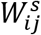 connects the neuron ‘j’ of the hidden layer to the output neuron, ‘i’. The bias of the hidden layer neurons and output neurons are represented by 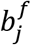 and 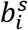 respectively.

The weights connecting the input layer and the hidden layer (*W^f^*) are absolute normalized.

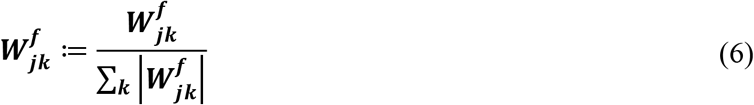

The bias of the neurons in the hidden layer depends on the energy released to it from the leaf node with which it is associated. The bias-energy relationship (Fig.2) is defined by equation (**7**). The higher the energy, the lower the bias, and hence higher the probability of firing for the neuron.

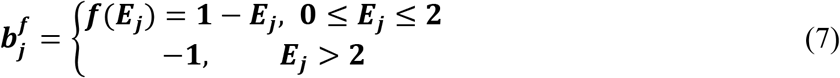

**Figure 2:**
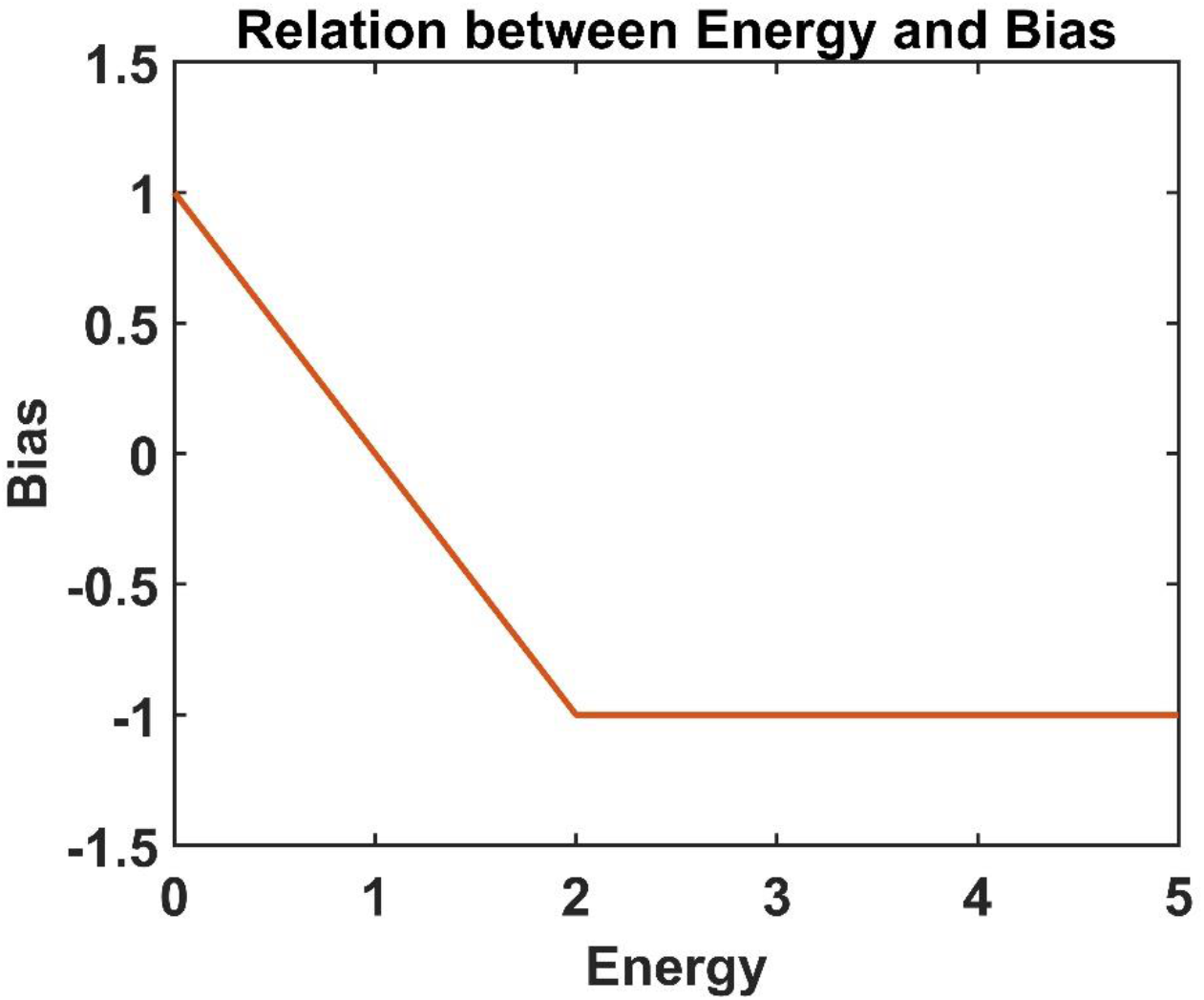
The relation between the bias of a neuron and the energy available at the associated leaf node

Given that the input vector is *x,* the net input to the hidden layer is given by

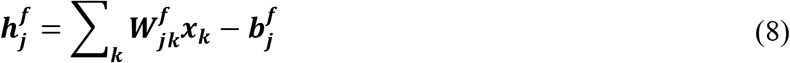

The output of the hidden neuron layer is obtained by passing the net input 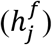 through a sigmoid function (*g*)

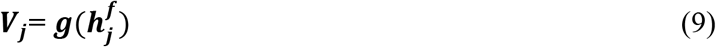

The net input to the output neurons hence can be written as

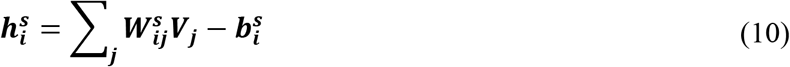

The output of each neuron is obtained by passing the net input through the sigmoid function

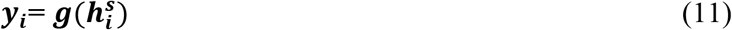

The weights and biases are updated to minimize the network error. Given that *d* is the desired output, the cost function without the regularization of energy is given by

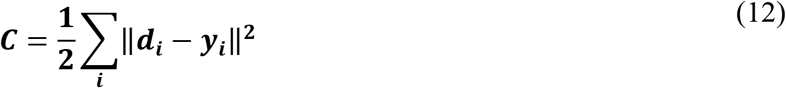

The gradients of the weights and biases are obtained as shown below in order to minimize the cost function

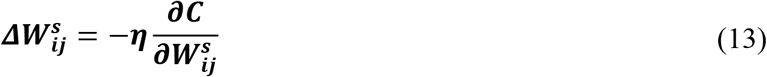

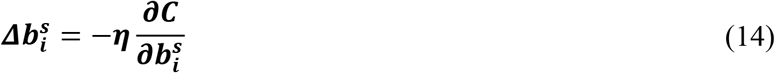

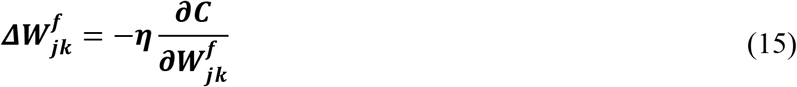

### Neurovascular coupling

Neurovascular coupling is implemented in this network in the form of dependence of the bias of each neuron to the energy level of the closest vascular node. Based on conductance-based neuron models, earlier studies have suggested that the effect of energy supply to a neuron can be expressed in the form of changes in the threshold of firing ^45,47^. During the forward pass, the bias of the hidden neurons is entirely dependent on the energy available at the leaf nodes (eqn. **7**). As shown in fig. 1, the state of the vascular network hence influences the activity of the neuron. The gradients at each level are calculated using the backpropagated error estimated at that level. Since the bias of the hidden layer is dependent on the leaf node energy, *E_j_*, available to that neuron, the gradient of the bias at the hidden neuron layer is converted in terms of the gradient of the energy as shown below.

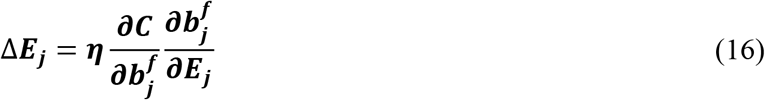

Given that the output error, *e_i_* = *d_i_* – *y_i_* the error terms at the hidden layer and output layer, 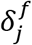 and 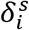 respectively are defined as below

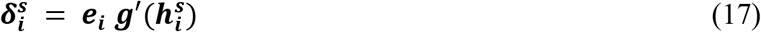

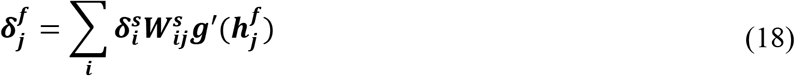

The gradients in terms of the partial derivative of the cost function can be rewritten using these error terms, as shown below.

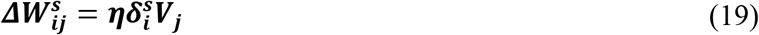

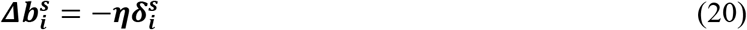

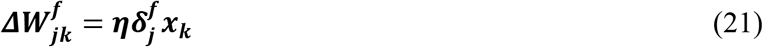

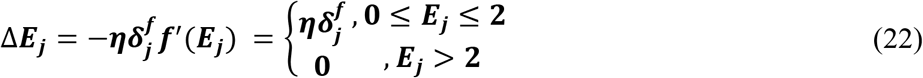

The gradient of energy calculated using eqn 22 is used to update the weights between a parent node *x* and child node *y* of the vascular tree.

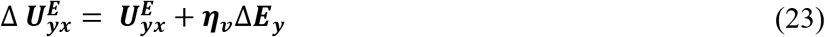

The energy gradient (Δ*E*_x_) at each parent node, *x* is taken as the average of the energy gradients of the children nodes.

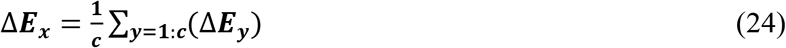

where c is the total number of child nodes of the parent node *x*.

In order to incorporate L^2^ regularization of weights, the cost function needs to be changed, as shown below. This, in turn, changes the gradient of *W_ij_*

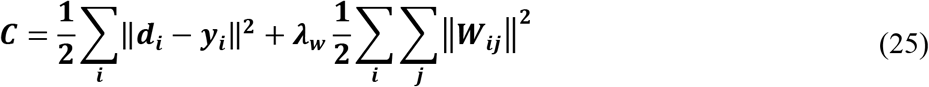

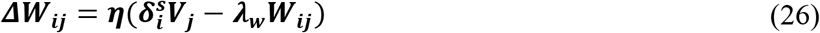

In order to incorporate L^1^ regularization of energy, the cost function needs to be changed, as shown below. This, in turn, changes the gradient of *E_j_*

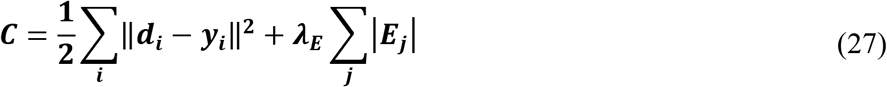

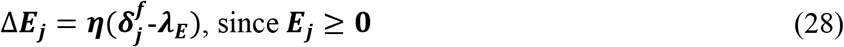

The performance of the ANVN is evaluated based on the accuracy (*α*) and the total energy consumed (ξ). The accuracy of the network (*α*) at any given root energy *ξ*, is defined as the fraction of the number of correct class predictions to the total number of predictions made from a given test data set.

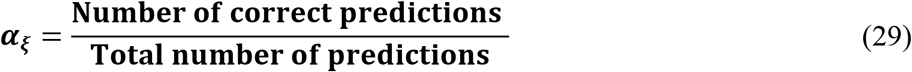

The total energy consumed (ξ) is calculated as the sum of the energies available at all the F leaf nodes, which subsequently is equal to the energy taken by the root node.

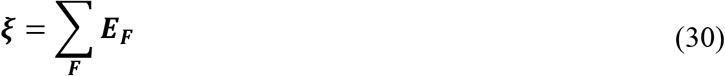

The minimum accuracy (*α*_*ξ*0_) of the network is the accuracy at minimum root energy (*ξ* = 1). The efficiency (*ψ_ξ_*) of the network at any root energy *ξ*, is defined as the ratio of relative accuracy and energy consumed.

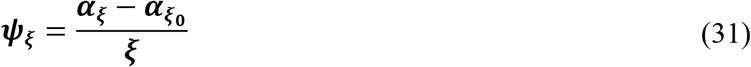

### Training regimes in the vascular network

In order to investigate the necessity of training the vascular network, we propose three training regimes to analyze ANVN. In all three regimes, the networks are trained using MNIST data set ^48^

### Untrained Vascular Network

Under this regime, the neural network is trained a priori without any vascular network. This trained network is then connected with the leaf nodes of the vascular network. The weights of the vascular tree are non-trainable and predefined such that the energy at the root node is equally distributed among the leaf nodes. The bias of the neurons now depends on the energy available at its leaf node. The bias derived from the energy at each neuron is calculated as per the relationship (eqn.**7**) shown in fig2. This untrained variant of ANVN is then tested using the same data set used to evaluate the performance of the trained MLP. The performance in terms of accuracy and efficiency is evaluated by varying the energy provided at the root node.

### Sequentially Trained Vascular Network

The second regime aims to study if training of the vascular network would lead to any improvement in the performance of the neural network by achieving an optimal delivery of energy to the hidden neurons. For testing this, the neural network is trained independently first. It is then incorporated in ANVN by connecting the hidden neurons to the leaf nodes of the vascular tree, and the weights of the tree are trained subsequently. The bias of the hidden neuron depends on the energy at the leaf nodes. This energy-dependent bias 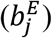 will be different from the trained bias 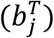 and the difference 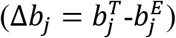 is used to calculate the energy gradient (Δ*E_j_*) using the bias energy relationship (eqn. **7**). This gradient of energy is used as vascular feedback to update the vascular weights.

### Simultaneously Trained Vascular Network

In this case, the neural and vascular networks are trained simultaneously. The untrained vascular network is connected to the hidden neurons of an untrained MLP to form the ANVN like before (fig(1)). During the forward pass of the MLP, the energy at the leaf nodes decides the bias of the hidden layer. The gradient in bias obtained during the backpropagation is used to find the gradient in energy (eqn.**16**). A neuronal update that seeks to reduce bias must demand more energy from the vascular tree. The energy gradient is an estimate of the neuron’ s energy deficit to attain the required change in bias. This information is propagated along the vascular tree, upwards from the leaf nodes to the root node, to modify the weights so that the energy supply at the leaf nodes match the neuronal demand.

### Simultaneously trained ANVN with an energy reservoir (ANVN_R)

In all three regimes discussed above, a pre-determined amount of energy is provided at the root node and distributed to all the hidden neurons by the leaf nodes. The network uses up the energy given to it regardless of whether the energy is in excess compared to their actual demand. Here we modify the ANVN such that the vascular network takes only the required energy from the root node, rejects the excess energy, and saves it in a reservoir. The ANVN with reservoir is denoted by ANVN_R. A modification was made to the calculation of energy gradient such that any neuron receiving a per capita energy > 2 units returns the excess energy by updating the weights using a small negative slope (γ=0.005). The energy gradient equation (**22**) was hence updated as shown below.

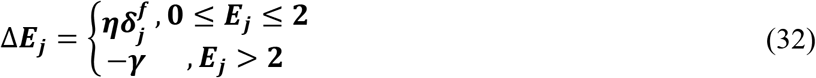

This energy gradient modifies the vascular weights from the leaf nodes to the root and also the weight connecting the root node to the source (fig. 1). Updating the source weight will naturally change the weight of the reservoir due to normalization carried out among vascular weights emerging from the same parent node (eqn. **3**).

## Results

### Performance of ANVN under various vascular training schemes

The initial part of the work focuses on establishing the importance of vascular training on the neural network. The ANVN was trained under 3 different regimes, and their performances were evaluated based on accuracy and energy efficiency.

#### Regime 1

Pre-trained neural network connected to untrained vascular tree

A neural network with 512 hidden neurons was trained independently using the MNIST dataset with 500 data points. The hidden layer of the neural network was then connected to an untrained vascular network with 512 leaf nodes. The branching factor of the tree was also varied such that the tree has a minimum of two levels (k=512) and a maximum of 10 levels (k=2). The result shown here is for k=8, which results in a tree with 4 levels. The bias of the hidden layer of the ANVN was now dependent on the energy available at the leaf nodes. The energy served at the root node was varied from 10 to 500 units. The performance of the ANVN was evaluated using test data set of 200 data point for each value of root energy over the same test data set used to test the performance of the independently trained neural network. The performance in terms of test accuracy and efficiency was abysmal for all root energies (dotted line with star marker in figFigure 3).

**Figure 3.**
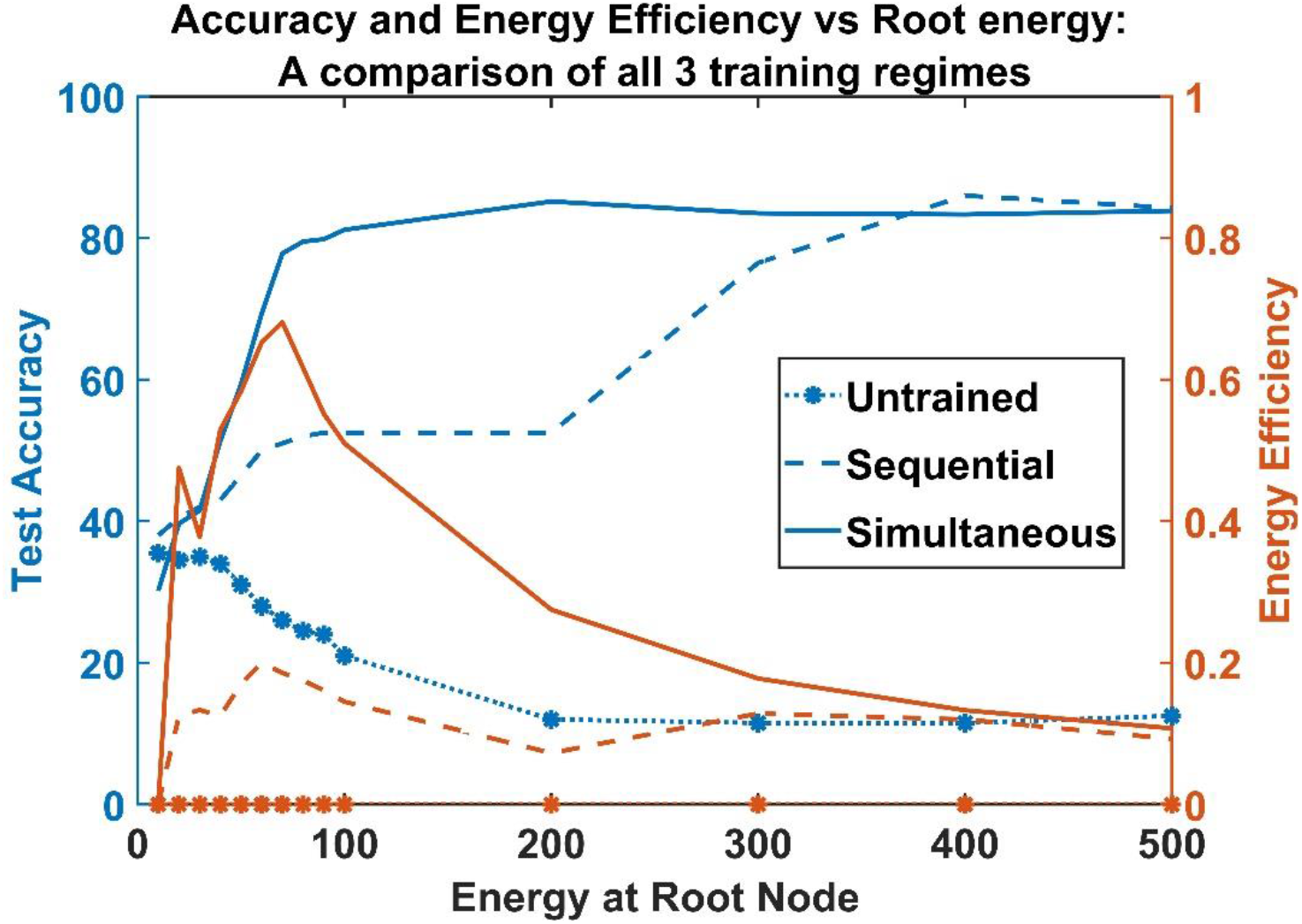
Accuracy and energy efficiency vs. root energy when vascular network is (i)Untrained (dotted line with star marker), (ii) Sequentially trained(dashed), and (iii) Simultaneously trained (solid) with MLP

#### Regime 2

Vascular tree sequentially trained after neural network training

The second training regime was to sequentially train the vascular network. The neural network was pretrained similar to that in regime 1. The trained neural network was then connected to a trainable vascular tree made of 512 leaf nodes with random weight initialization. The difference in the trained bias and the bias obtained from the energy at the leaf node was used to train the vascular network. The vascular network was also trained using the same MNIST data set (500 training data points) used to train the neural network. Once the vascular network was trained sequentially, the performance of ANVN was evaluated using 200 test data points by varying the energy supplied at the root node. The energy was varied from 10 units to 500 units, and the accuracy (eqn.**29**) and efficiency (eqn.**31**) were calculated in each case. The maximum efficiency attained by the network was just around 0.2 units. The network showed an improved accuracy at higher energies (> 300 units) (fig.Figure 3 blue dashed line). But the efficiency plot shows a steady decrease with an increase in root energy.

#### Regime 3

Simultaneously trained MLP and Vascular tree

The third training regime followed was training both the MLP and the vascular network simultaneously. The network is trained using 500 data points of the MNIST data set. The gradient of bias obtained using the backpropagation algorithm was converted into a gradient of energy in order to update the vascular weights. The training was carried out for 20k epochs by varying the root energy from 10 to 500 units. The network was then evaluated using the test data set of 200 data points. The simultaneous training of the vascular network and neural network improved the network performance even further. As shown by the solid line in fig. 3, the network attained a peak efficiency of 0.62 units with test accuracy above 80% at much lesser energy (100 units) compared to the networks trained using regimes 1 and 2. The network was able to achieve better efficiency at lower energy range, and on further increase in root energy, even though accuracy was maintained, the efficiency kept dropping steadily. Thus, simultaneous training of neural and vascular networks is desirable to carry out energy-efficient data processing.

### Network performance is invariant with respect to the branching factor

The vascular tree in this model is determined by the number of hidden neurons of MLP, (N) (which equals the number of leaf nodes), and the branching factor (k). Change in the branching factor changes the topology of the tree. We now consider the effect of the branching factor, k, on MLP learning.

Let the total number of vascular nodes be *t_n_* and vascular nodes *x* and *y* be connected by weights 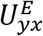. The vascular root node is supplied by an energy source, *E_s_*. The parameters of the model comprise the values of the weights and input energy. In the described tree, every node except the root node is connected to its parent node; hence there exists *t_n_* – 1 weight connections. Including the input energy (root node energy), the total number of parameters add up to (*t_n_* – 1) + 1 = *t_n_*.

The normalization of weights at each node forms the constraint of the model along with the energy consumed at each vascular node. Since the number of weight connections is *t_n_* – *N*, the number of constraints resulting from the normalization of weights account to *t_n_* – *N*. The second set of constraints are the energy consumed at each vascular leaf node. Since there are N leaf nodes, the total number of constraints of the network adds up to (*t_n_* – *N*) + *N* = *t_n_*.

Hence regardless of the branching factor, the number of parameters = the number of constraints, making the solution unique. This feature was observed in the model from the similar network characteristics exhibited by networks trained using all the three training paradigms (untrained neural, sequentially trained, and simultaneously trained) regardless of the variation in the branching factor, k (fig.4, fig.5 and fig.6).

**Figure 4.**
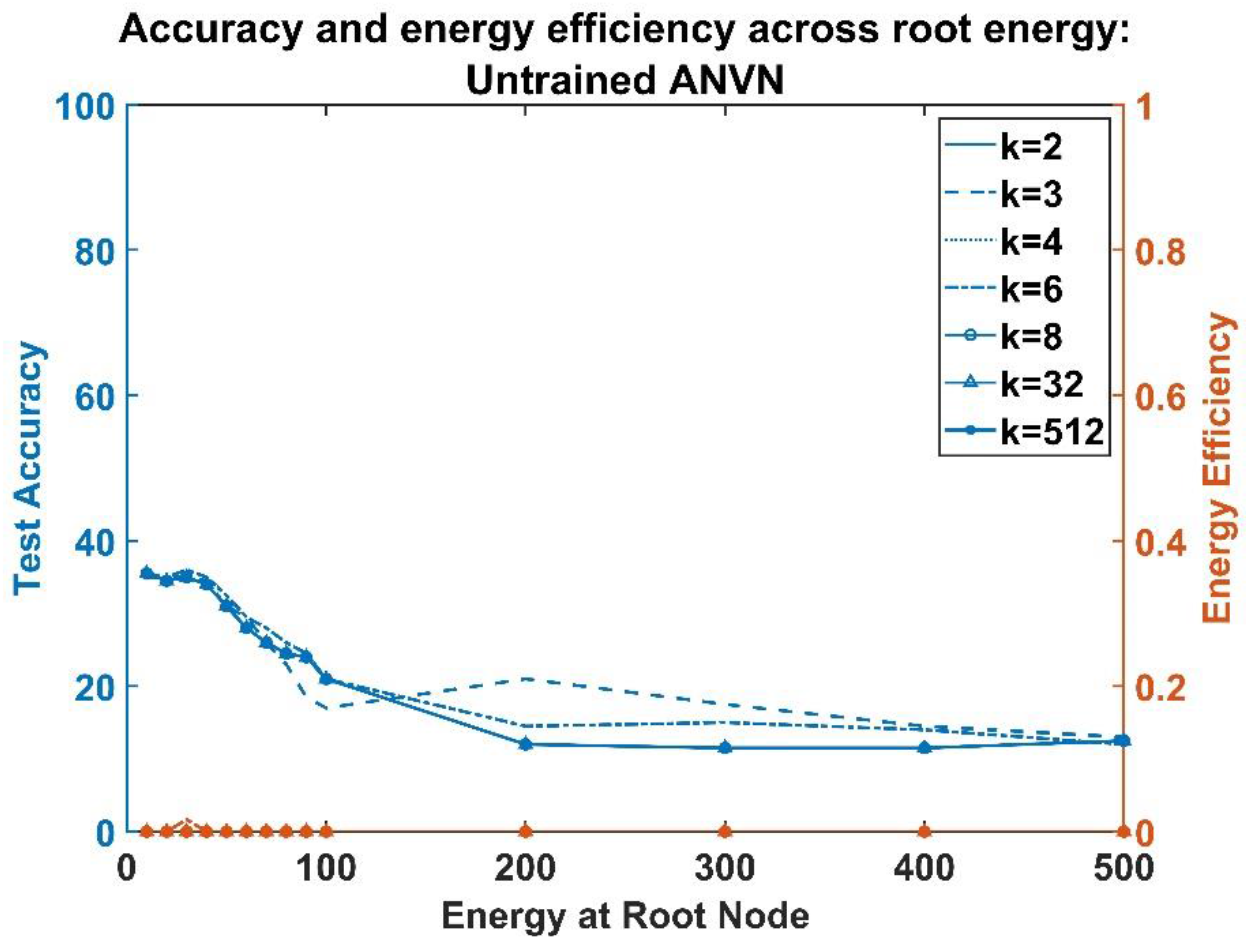
Untrained ANVN: Energy Efficiency and accuracy across root node for various branching factors

**Figure 5.**
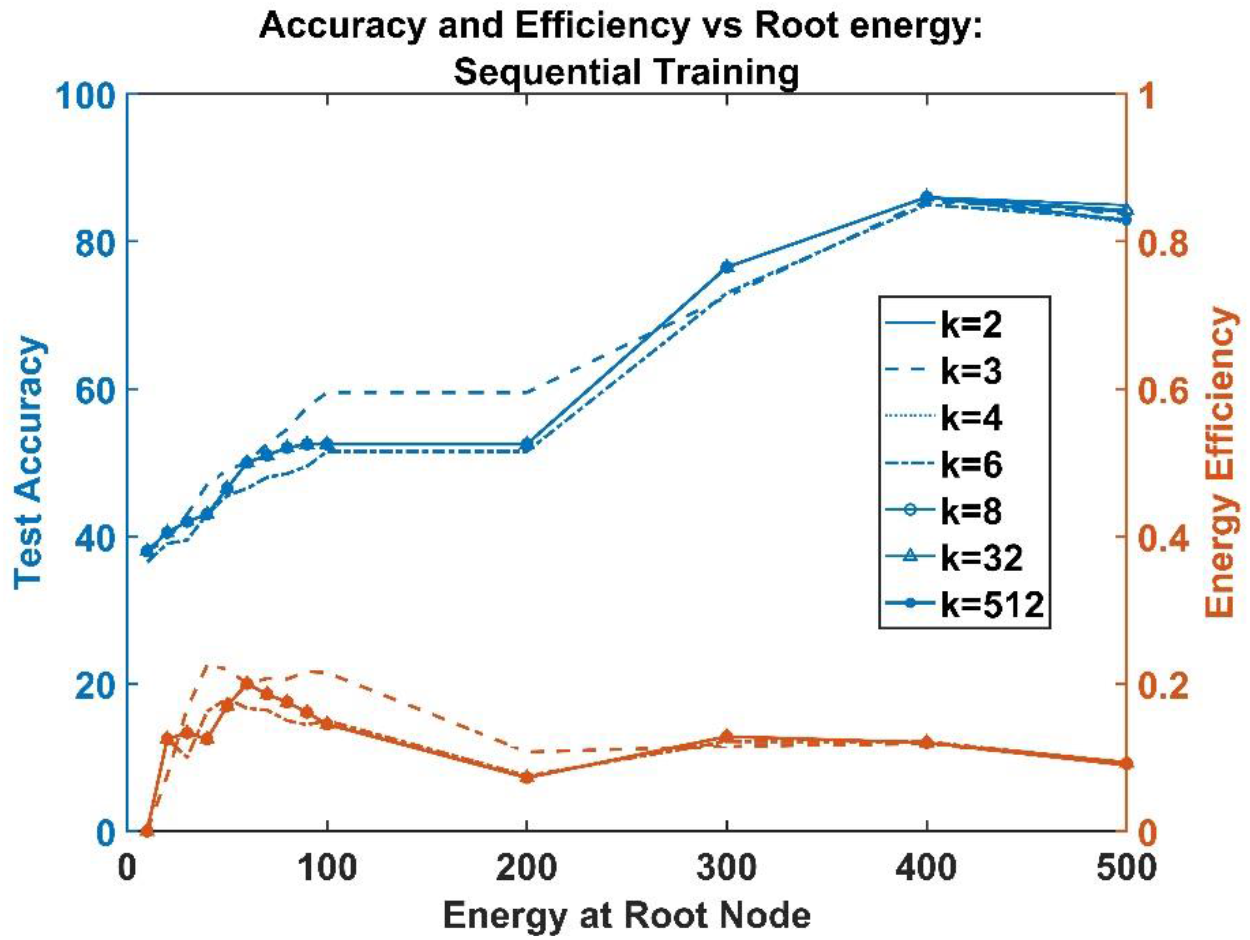
Sequentially trained ANVN: Energy Efficiency and accuracy across root node for various branching factors

**Figure 6.**
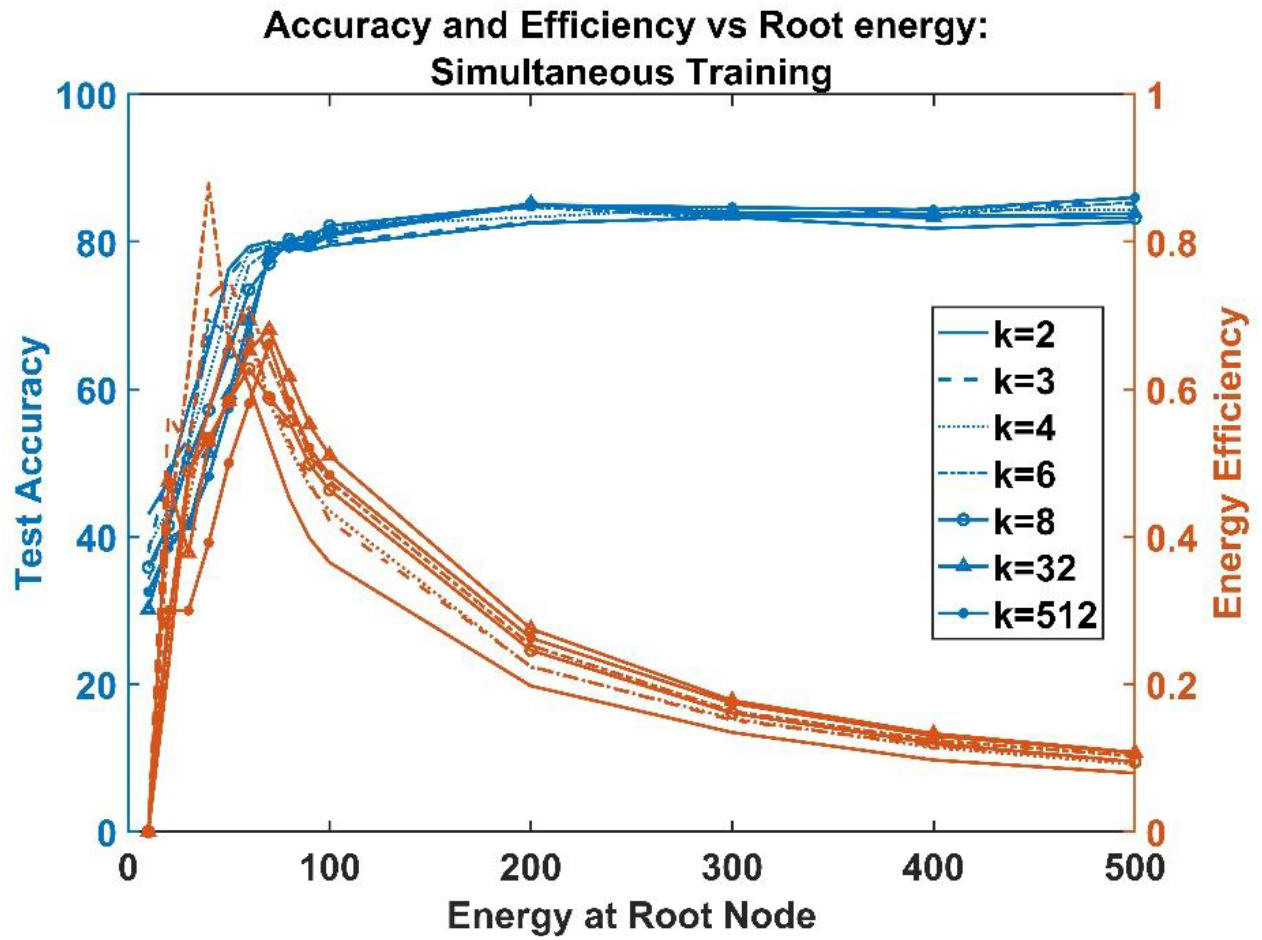
Simultaneously trained ANVN: Energy Efficiency and accuracy across root node for various branching factors

### Energy deficit vs. accuracy correlation study for various root energies

To study the effect of energy on the accuracy of the network, we need to check if the energy deficit would result in a drop in the accuracy. We define a term, ‘energy deficit’ 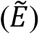 as the difference of desired energy (*E_D_*) and actual available energy (*E_A_*). The desired energy of a pretrained neural network is the total energy calculated using the trained biases 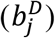 based on eqn (**7**). The available energy at an ANVN is the sum of energies at the leaf node.

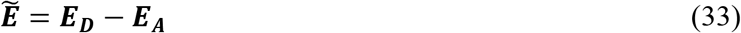

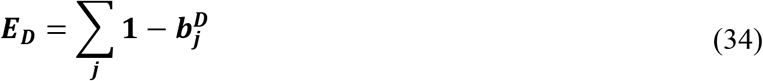

It is easier to quantify energy deficit using the first and second training schemes (untrained and sequential training). For a sequentially trained ANVN, the vascular tree is trained after neural network training to match the desired energy to provide a bias close to the trained neural network. Here there is a provision to cross-check if there is an actual deficit in energy by calculating the deviation of the trained vascular leaf node energy from the desired energy. The variation of energy deficit (*E_D_*) and accuracy (*α*) as a function of root energy (*E_s_*) is plotted in fig (7.a). The network has 512 neurons in the hidden layer, and the branching factor of the vascular tree is 32. The training and testing procedures are similar to the sequential training procedure described under the subsection, ‘Regime 2’ . At lower root energies, the energy deficit is high, and the accuracy is low. As the source energy increases, the energy deficit reduces, and accuracy improves. Estimation of correlation between Energy Deficit 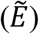 and Accuracy (*α*) showed a strong negative correlation (Pearson correlation coefficient=-0.94). This showed that energy deficit inversely affects the accuracy of the network.

**Figure 7.**
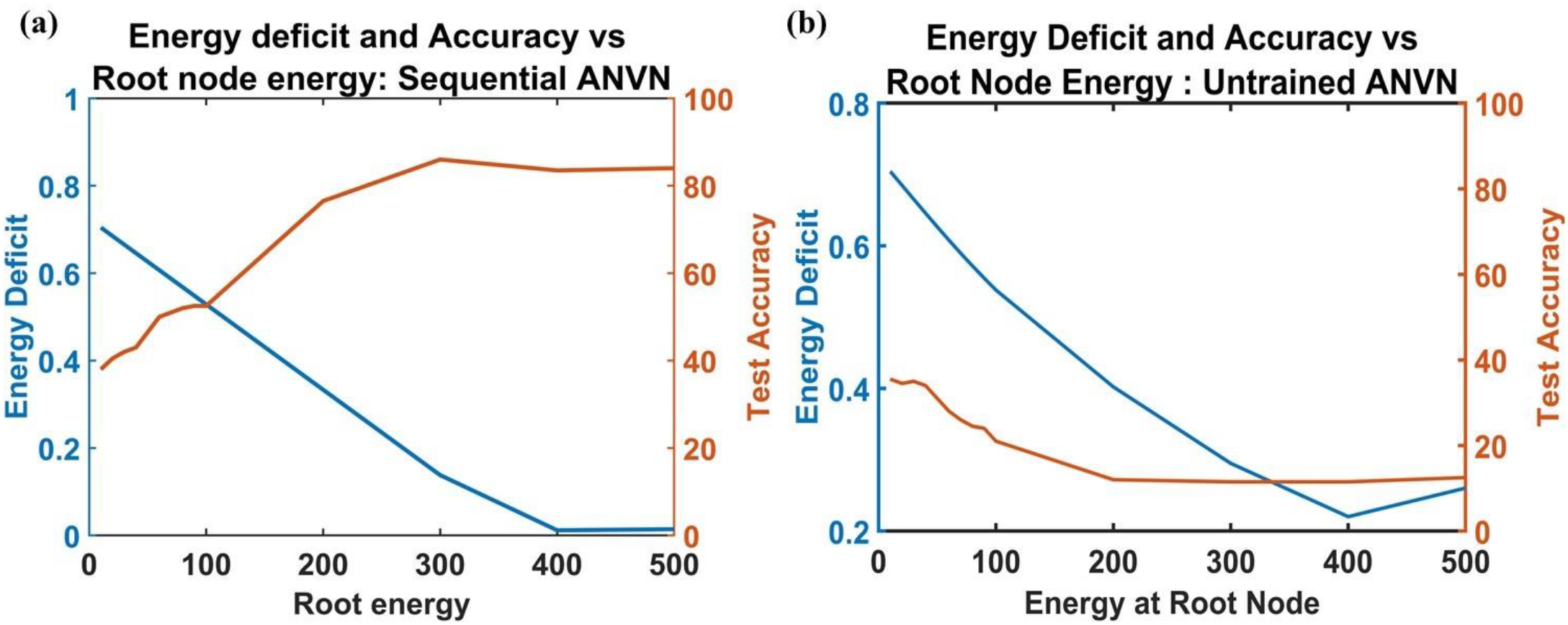
Accuracy and energy deficit variation across root energy (a) for sequentially trained ANVN (b) for untrained ANVN

In untrained ANVN, the desired energy of the network is unknown to the vascular network due to a lack of vascular training. Hence, the accuracy shows no improvement (fig 7.b) even when the energy deficit is low, once again highlighting the point that high energy input is not of any advantage unless the vascular network is trained. The network size and branching factor are the same as the sequential ANVN.

### Simultaneously trained ANVN with an energy reservoir

A comparison of vascular training regimes 1,2, and 3 established that the network performance tremendously improves at lower source energy when the vascular network is simultaneously trained with the neural network. The simultaneously trained ANVN attained high accuracy at relatively smaller root energy, but the accuracy did not improve on further increase in root energy as observed in figFigure 3. This was reflected in the steady decrease in efficiency of the network. An ideal network should be able to reject the unwanted excess energy provided to it. The ANVN was hence modified to obtain the energy from the source as demanded by the neural network and reject the rest so that it can be saved in the reservoir.

The root node of the network was connected to a constant energy source of value 5000 units through a weighted connection (fig.1). Another weighted connection from the energy source connected it to the reservoir. If the weight of the connection to the root node of ANVN_R is defined as *U*_0_, then the weight connecting the energy source and reservoir would become 1 – *U*_0_. The vascular weights, including *U* were trained so that only the energy demanded by the network would be received, and any excess energy (i.e., energy not taken up by the neural network) would be pushed into the reservoir. The initial value of *U*_0_ determines the initial energy available to the hidden neurons. The total energy consumed by the neural network depends on the demand of the neurons in the hidden layer. Hence limiting the number of neurons in the hidden layer would become critical. The ANVN_R was hence studied by varying the number of hidden neurons.

The performance of the ANVN_R was evaluated in terms of accuracy and efficiency. The efficiency (*ψ_n_*) of a network with ‘n’ number of hidden neurons (1 < *n* < *N*) in the hidden layer was defined as the fraction of relative accuracy and energy consumed by the network. The relative accuracy was calculated as the difference of accuracy using any given number of hidden neurons (*α_n_*) and the accuracy achieved using the minimum number of neurons (*α*_*n*0_), *n* = *n*_0_. In the current model, the minimum number of hidden neurons used was *n*_0_ = 16. The total energy consumed (*ξ_n_*) was calculated by the sum of the energies available at the terminal leaf nodes of the vascular tree.

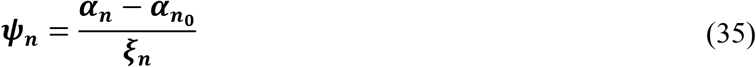

The network was observed by varying the number of neurons in the hidden layer between 16 and 500. Each network was probed multiple times by varying the initial average per capita energy received by the leaf nodes between 0.2 units and 1 unit by adjusting the initial value of *U*_0_. Similar to the ANVN, we used 500 data points for training and 200 data points for testing the network. The training was carried out for 20k epochs.

Figures (8a,9) show that the accuracy of the network increased sharply (slope~1.4) with an increase in the number of hidden neurons until N=32. Further increase in the number of hidden neurons resulted in a relatively slower improvement in accuracy (slope~0.36) till N=64. The increase in accuracy became even slower beyond N=64 (slope~0.04) till N=175, and a further increase in the number of hidden neurons almost saturated the gain in accuracy. Nevertheless, it is interesting to note that the highest accuracy (98%) attained by the ANVN_R (N=500) was much higher than the accuracy obtained by simultaneous training of ANVN (N=512) without reservoir (85%). The efficiency of the network peaked when N was in the range 28 to 36 (fig. 8a) and then fell systematically with a further increase in the number of hidden neurons. Even though the accuracy increased from approximately 80% at N~36 to around 98% at N~500, the efficiency started to drop beyond N~36. This shows that for the network to be maximally efficient, it needs to compromise the maximum accuracy attainable. Even though the average per capita energy consumption showed a slightly decreasing trend with an increase in the number of hidden neurons (fig.9.a), the total energy consumption increased linearly (fig.9b), which in turn resulted in the fall in efficiency.

**Figure 8.**
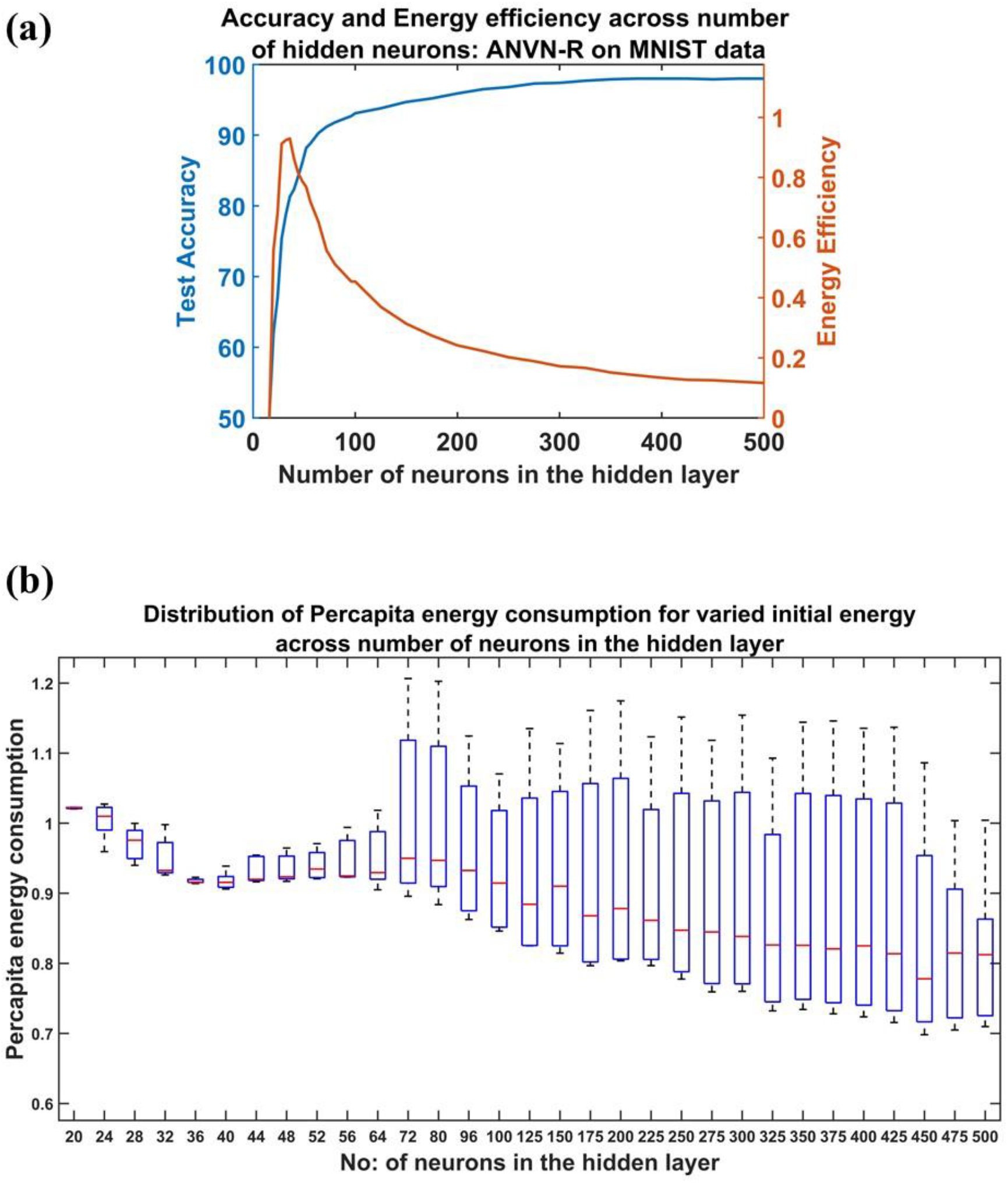
Study of ANVN_R: (a) The test accuracy and energy efficiency across the number of hidden layer neurons. (b) The box plot for visualizing the settling points of per capita energy consumption for each initial energy (varied from 0.2 units to 1 unit) given to single neurons given for each network size. The red mark shows the median value of the per capita energy consumed by the trained network and its maximum and the minimum values determine the height of the box.

**Figure 9:**
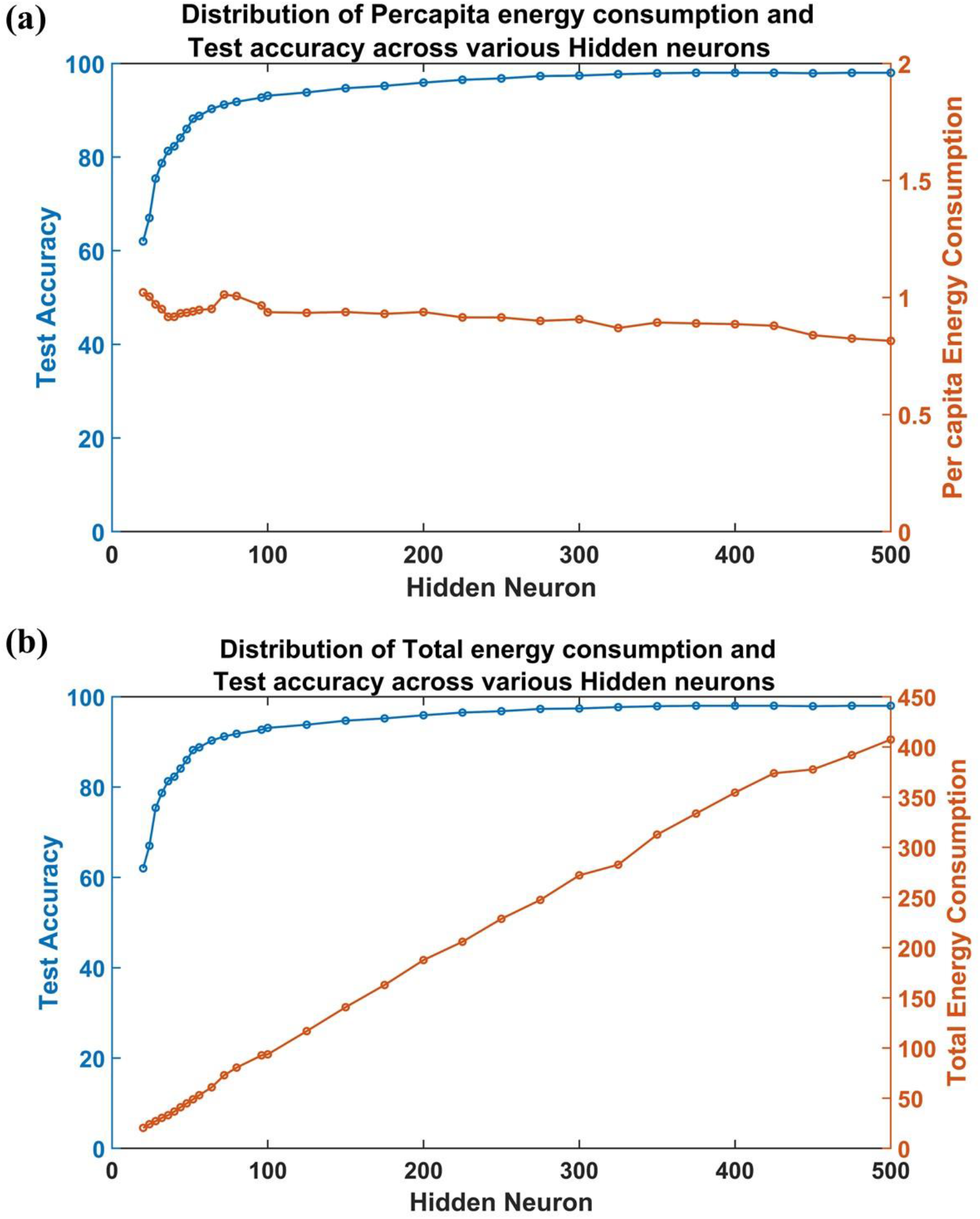
Study of variation in accuracy and energy consumption in ANVN_R with an increase in the number of hidden neurons: (a) Test accuracy vs. per capita energy consumption (b) Test accuracy vs. total energy consumption

Each network with a given number of hidden neurons was observed by varying the initial per capita energy made available to the neurons. This variation in the initial per capita energy, delivered by the leaf nodes of the vascular tree to the hidden neurons, resulted in an interesting observation. The pattern of variation of accuracy against energy consumption as epochs progressed differed for networks with different number of hidden neurons.

fig. 10 a,b,c shows the trajectory of the network evolution in the space of per capita energy consumption vs. accuracy for various initial conditions. As epochs progressed, the trajectories seemed to converge to a point (fig 10.a) when the number of hidden neurons was small (N<64). This point appears to be a stable fixed point of the network dynamics on the per capita energy vs. accuracy space. Further increase in the number of hidden neurons, the trajectories did not converge to a point anymore but seemed to approach a line. It appears the former fixed point had given place to a line of attractor as shown in fig 10.c. The transition can be observed clearly in the box plot shown in fig.8b. The box plots describe the variation in per capita energy consumption of the networks at the steady state given a range of initial per capita energies (0.2 units to 1unit). By steady state, we mean that the network is trained for a sufficiently long time (20k epochs). The red mark in each box shows the median value of steady state per capita energy consumption across varied initial per capita energy of a network with a given number of hidden neurons (N). The maximum and the minimum values of the per capita energy consumed by the trained network determines the height of the box. The height of the box was small for smaller networks (N<64), which indicated a low variation in energy consumption, indicating a fixed point attractor. As the number of hidden neurons was increased, the height of the box increased, indicating a line of attractor for steady state per capita energy consumption. The transition (fig 10.b) of the network stable state from a fixed point to a line of attractors happened between the hidden neurons number 44 to 64 (fig.8.b).

**Figure 10.**
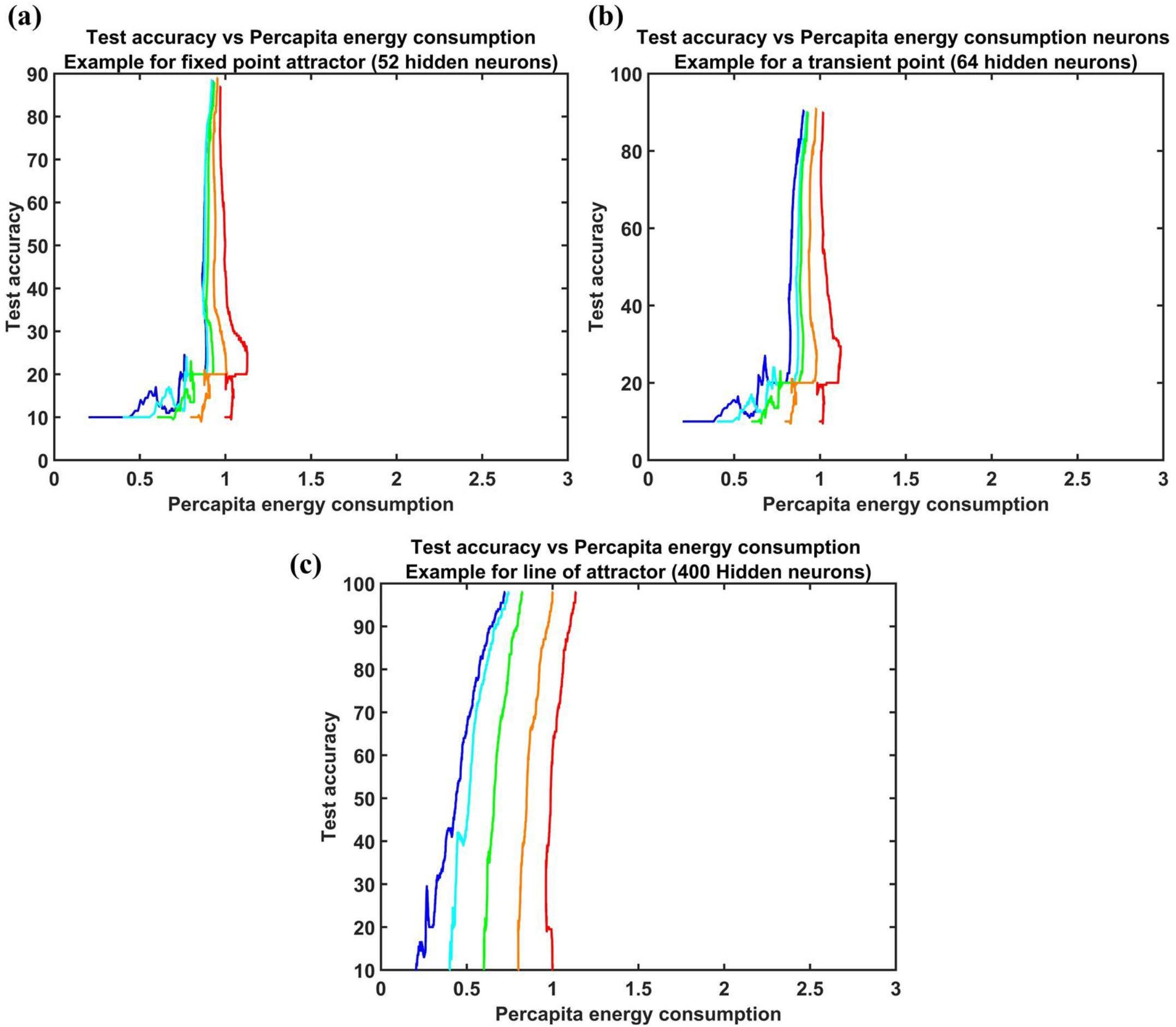
Study of variation in accuracy and energy consumption with an increase in the number of hidden neurons. Each color indicated the trajectory of individual simulations with different initial energy given to the network. The starting point at the x axis denotes the initial energy of each simulations: (a) An example of the trajectories converging to fixed-point attractor (b) An example of a transition point from fixed point to line of attractors (c) An example where trajectories converge to a line of attractor.

To summarize, for smaller networks, the network seeks the same point in the per capita energy vs. accuracy space irrespective of the initial conditions. On the other hand, for larger networks, the final state of the network was strongly dependent on the initial per capita energy. Furthermore, in the latter case, the final state shows variation primarily in the per capita energy and not in the final accuracy achieved. It is also interesting to note that this transition happens roughly at the same number of hidden neurons even when the network is trained with a different data set (EMNIST - results in the supplementary material fig S1, fig S2).

### Transfer learning

The importance of vascular training in ensuring optimal performance of the neural network was proven by the results discussed earlier in this study. The plasticity of vasculature is observed experimentally in many recent studies ^49–52^. This posed the following question: how effectively does a vascular tree trained on data set A meet the energy demands of the network when it is trained subsequently on data set B? In order to answer this question, the simultaneously trained ANVN_R was initially trained with MNIST data^48^, and then the vascular weights were frozen to use as the initial weights for training a different data set (EMNIST^53^ data in this case). The neural network of ANVN_R had 100 neurons in the hidden layer, and the vascular network was assigned a branching factor of k=8 in order to explore the changes in vascular weights in 4 levels of branching. Training using the MNIST data set was carried out for 20k epochs with 500 training samples and was tested using 200 data points. Similarly, the training using the EMNIST data set also was carried out for 20k epochs using 500 training data points. The EMNIST data (200 points) were used to test the trained network. The difference in the vascular weights (*U_A_*) of ANVN_R trained using MNIST and the weights (*U_B_*) of ANVN_R trained using EMNIST was quantified using the Root Mean Squared Error (RMSE) between them. Each vascular node was assigned a level number (1 < *l* < L) based on its hierarchical position from root node, with root node being at level, *l* = 1. The RMSE between *U_A_* and *U_B_* was estimated for each level *l* by considering the weights emerging from all nodes (*i* ∈ *1*) in the same level.

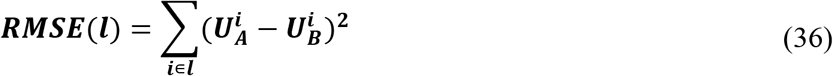

The ANVN_R that was already trained on MNIST data learned the EMNIST data set, which is similar to MNIST, much faster, as shown in fig (11). The red curve indicating the training accuracy of the network trained on EMNIST increased to approximately 80% very quickly. In fig (12a), the RMSE in each level starting from the level of root node (Level 1) to leaf nodes (Level 4) is plotted. The variation of RMSE with respect to levels in the vascular tree showed that the most significant changes happened at level 4, which is the level of the leaf nodes and hence closest to the neurons. The RMSE between *U_A_* and *U_B_* systematically reduced from level 4 to level 1, which means that the maximum changes occurred at the leaf node, and the changes were minimal at levels farthest from the leaf nodes and close to the root node. As the training progressed, this difference in the vascular weights at various levels in the vascular tree became more prominent (fig12.b). The change followed a similar pattern when explored in the network without reservoir (ANVN trained under regime 3 – simultaneous straining), simulated for a larger number of levels (L=7) (Fig. S3 and S4 in supplementary material).

**Figure 11.**
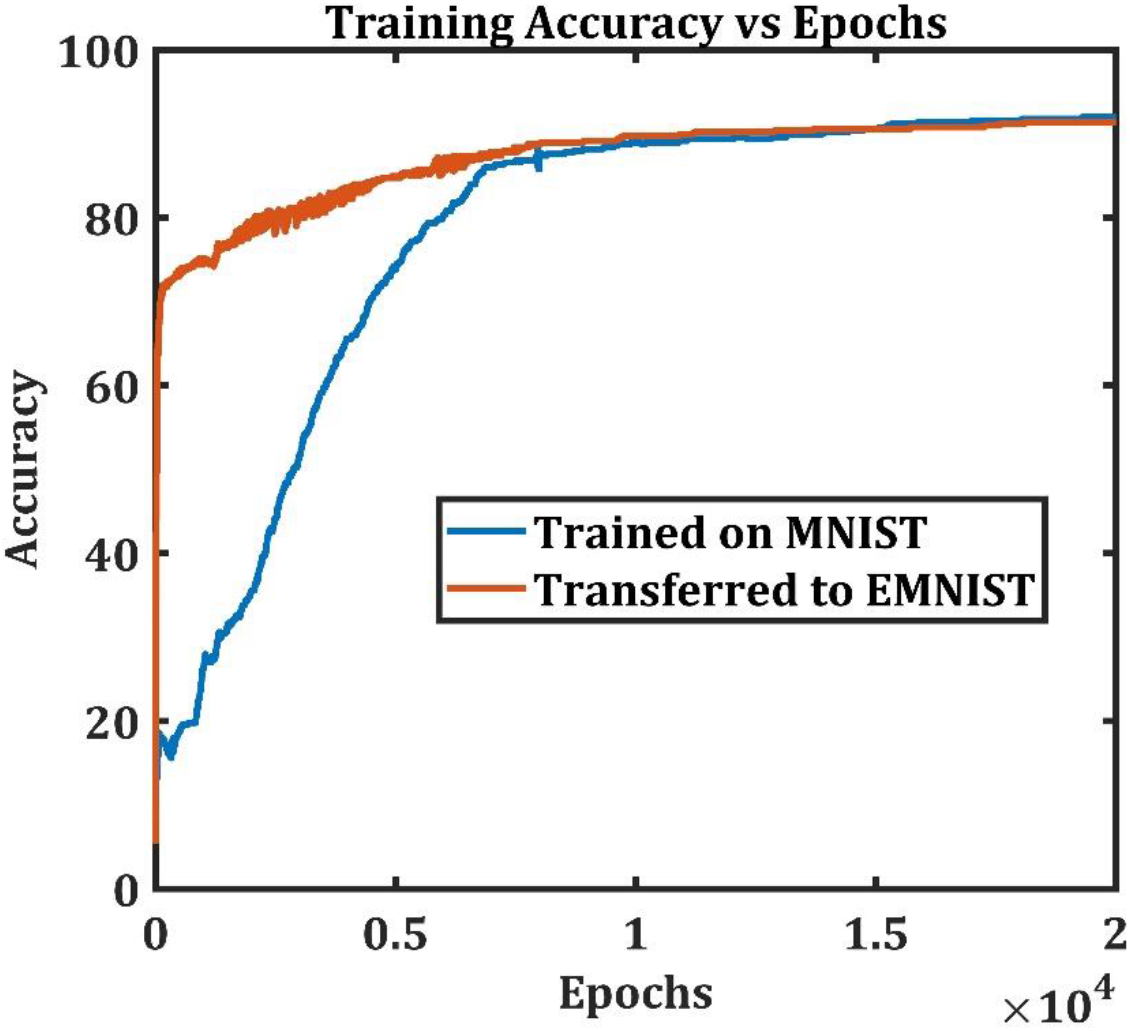
Training accuracy of the initially trained network (blue) takes more epochs to reach a high accuracy as compared to the network trained by transfer learning (red)

**Figure 12.**
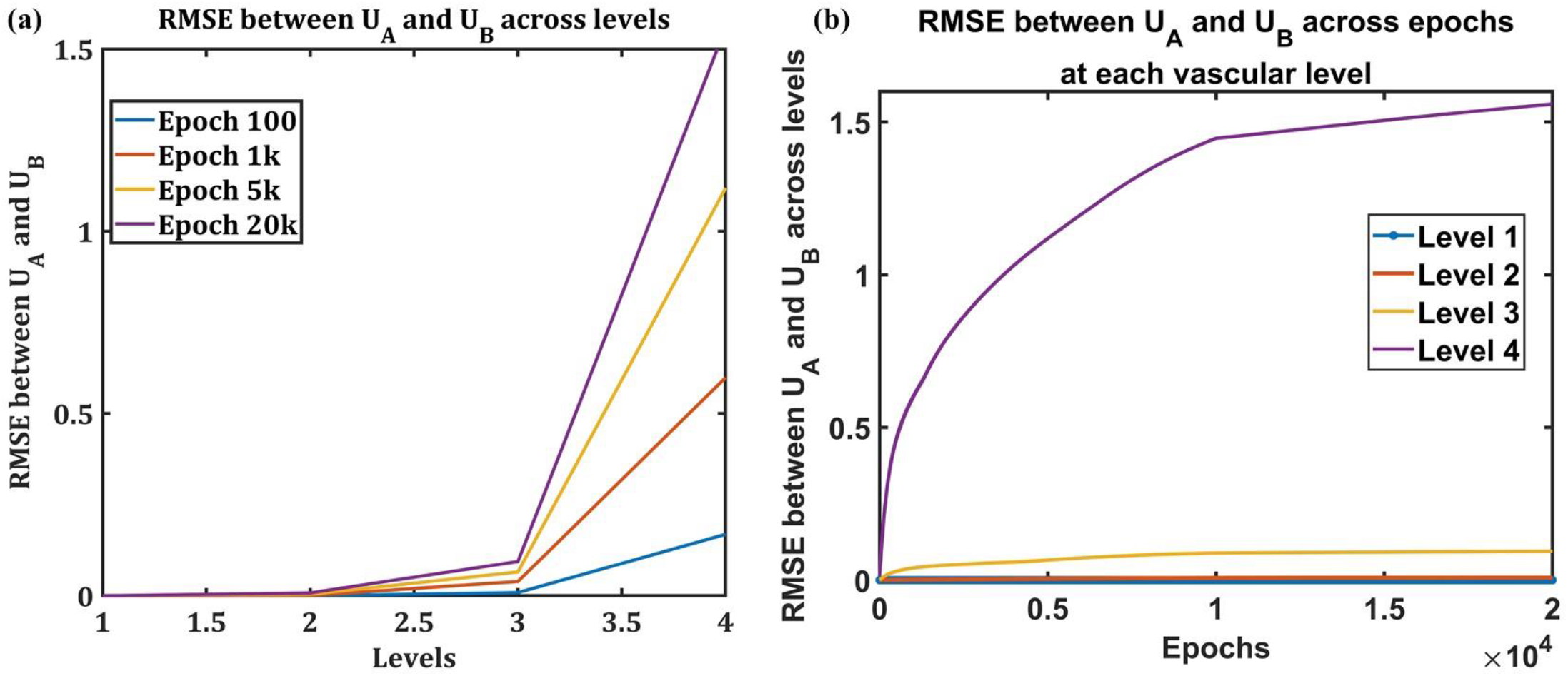
Variation of RMSE of the vascular weight at each level in the vascular tree (a) The RMSE between vascular weights at each level. The colors represent the RMSE at specified epochs. (b) The variation of RMSE across epochs for each level

### Regularization of Energy

The simultaneous training of neural and vascular networks in ANVN_R showed that increasing the number of hidden neurons beyond a point increases the total energy consumption without much improvement in accuracy. Such behavior might be a consequence of cost function not explicitly demanding minimal energy consumption. Hence, we decided to study the effects of constraining the cost function by introducing regularization. Regularization of weights is a technique widely used to improve the generalization of a neural network (chapter 3.4, Hastie et al., 2008). It prevents the overfitting of data. We explored if regularization of energy can bring about any change in the performance of ANVN. Two methods of regularization were explored. One was by penalizing the weights, by implementing *L*^2^ regularization of weights (eqn.**25**). The second method was directly constraining the energy by imposing *L*^1^ regularization of energy (eqn.**27**). For directly including regularization of energy in the cost function, the *L*^1^ regularization of was preferred over *L*^2^ since the biological implication of *L*^1^ minimization of energy was more meaningful. The range of initial values of average per capita energy was varied between 0.4 units and 1 unit. Any neuron receiving a per capita energy > 2 units (due to random initialization of the vascular weights) will return the excess energy by updating the weights using a small negative slope (eqn.**32**). The training data and testing were done using 500 and 200 data points, respectively, from the MNIST data set. Each network was trained for 20k epochs.

The *L*^2^ regularization of weights did not show any notable drop in energy consumption with an increase in the number of hidden neurons (fig. 13.b and 14.b). Instead, the accuracy dropped on introducing *L*^2^ regularization. However, on imposing the constraint directly on the magnitude of the energy consumed by the hidden neurons (*L*^1^ regularization of energy), a significant drop in energy consumption was observed with regularization (Fig 13.a and 14.a). The drop was higher with a higher regularization factor (λ). Moreover, the accuracy was maintained similar to that without regularization. Also, the transition of the network from a fixed-point attractor to a line of attractor appeared to occur much later in the case of a network regularized using *L*^1^ norm of energy (fig. 15.a) when compared to the non-regularized network (fig. 8) and the network regularized using *L*^2^ norm of weights (fig. 15.b). On imposing regularization of energy, the network converges to a fixed-point attractor in the per capita energy consumption vs. accuracy space for a range of larger networks making it more robust to variation in initial energy. Due to the large variation in the accuracy attained by the smallest network across the regularization factors, the relative accuracy, and hence by current definition, the efficiency also cannot be compared across the λ values.

**Figure 13:**
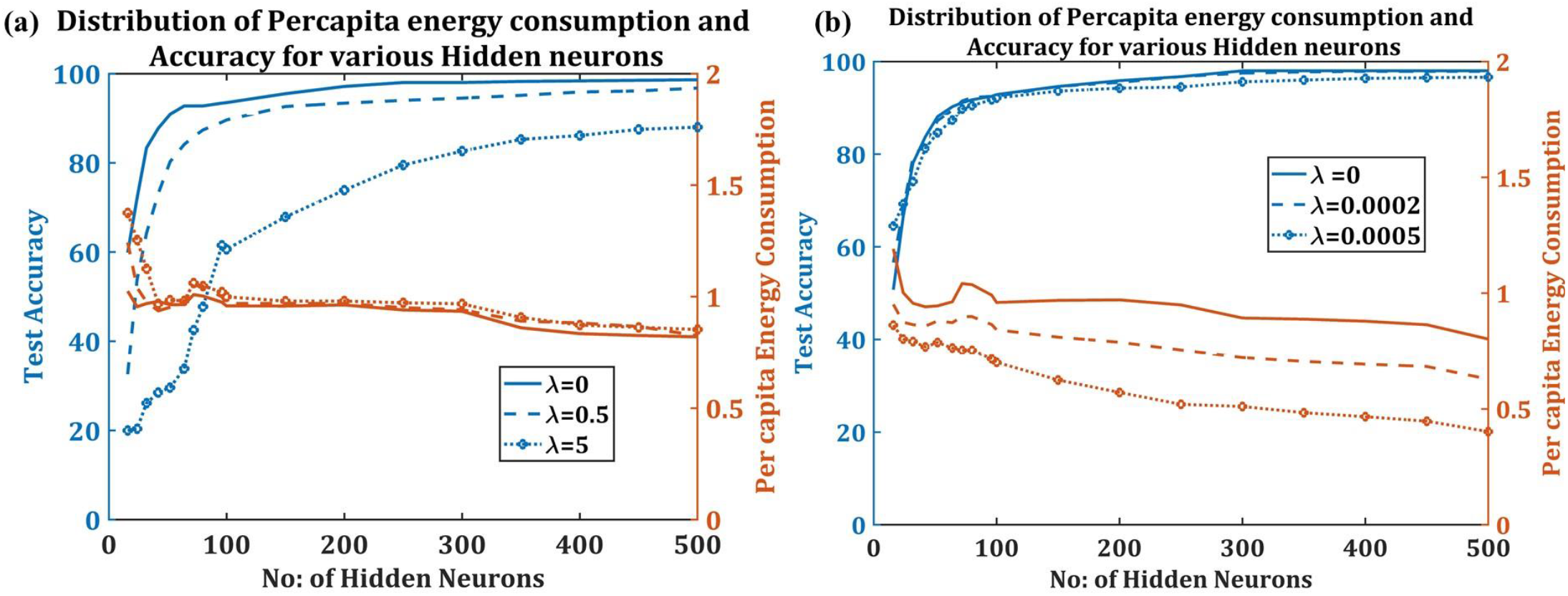
Comparison of performance (Test accuracy vs. Per capita energy consumption) at different values of regularization coefficient, *λ* in case of (a) *L*^1^ regularization of energy (b) *L*^2^ regularization of weights

**Figure 14:**
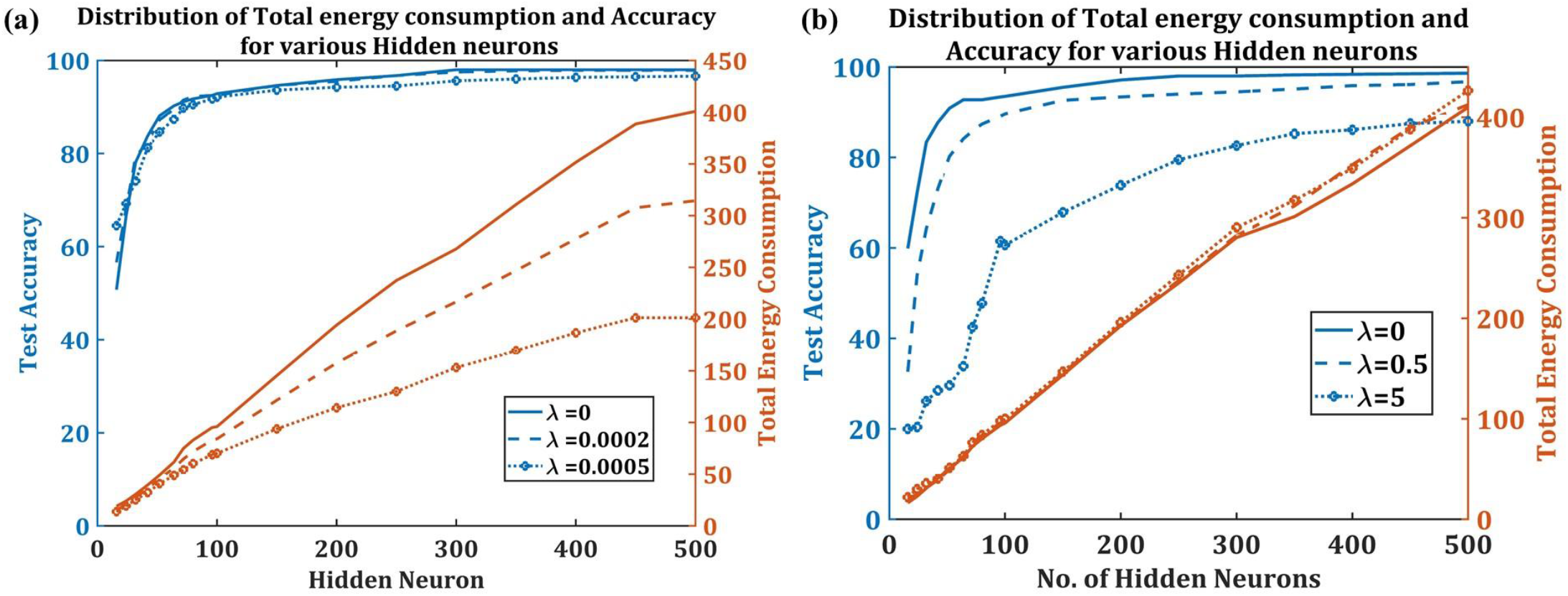
Comparison of performance (Test accuracy vs. Total Energy consumption) at different values of regularization coefficient, *λ* in case of (a) *L*^1^ regularization of energy (b) *L*^2^ regularization of weights

**Figure 15:**
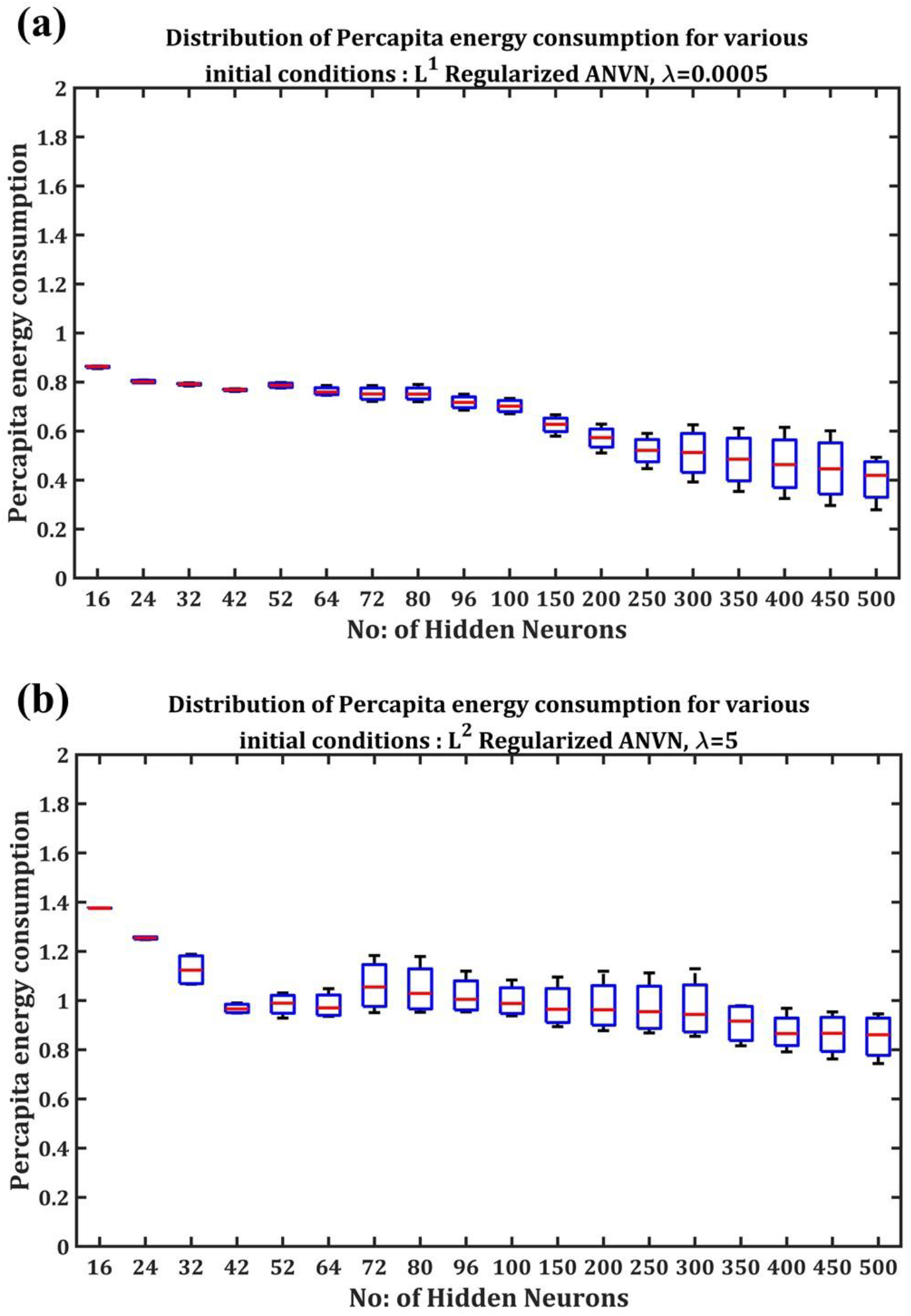
Box plot of variation in percapita energy at different initialization while incorporating (a) *L*^1^ regularization of Energy (b) *L*^2^ regularization of weights

### Correlation between error contribution and energy consumption

The hidden neurons in an ANVN and ANVN_R network consume, at the single neuron level, energy ranging from a minimum value of 0 units to a maximum of 2 in ANVN_R and more than that in the case of ANVN. A few neurons appeared to consume higher energy compared to other neurons in the same network. This variation of energy consumption among individual neurons of a network prompted the following question: Is the energy consumed by an individual neuron related to its contribution to the network’s performance? In order to answer this question, the correlation between the error contributed by each neuron and the energy consumed by the same neuron was calculated. To this end, during testing, for a network with N hidden neurons, neuron ‘j’ was switched off by making its output zero, and the test error (*ε_N_*(*j*)) was observed. The measured error is a readout of the contribution of the switched-off neuron to the network performance. Given a test sample size of M data points, the prediction error was calculated in terms of the root mean squared error (RMSE) between the desired output *d_i_* and the predicted output 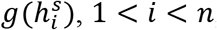, where *n* is the number of neurons in the output layer.

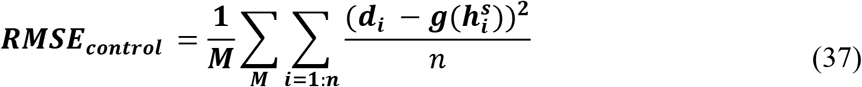

Since the data used was from the MNIST data set, the labels range from 0 to 9. Thus, for a given data point *p*, if the class number is *φ^p^*, the desired output *d* was defined such that

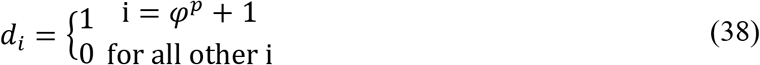

The output of the *j^th^* neuron in the hidden layer *V_j_* is obtained by passing the net input to the hidden layer 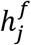 (eqn. **8**) through a sigmoid function (*g*) as explained earlier in eqn. **9**.

In order to calculate the error contribution, RMSE is estimated while switching off neurons 1 to j one by one for a network with j ranging from 1 to N. While switching of *k^th^* hidden neuron, the eqn. **9** describing the output of the hidden layer (***V_j_***) was modified

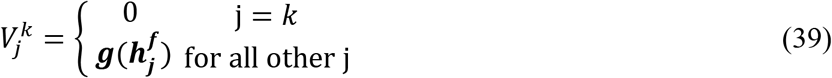

The root mean squared error obtained by shutting off *k^th^* neuron (RMSE_*k*_) was calculated as below

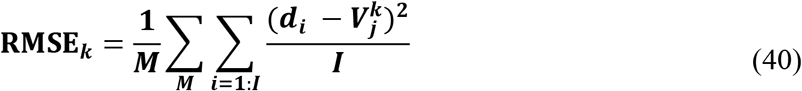

The gradient ΔRMSE for each hidden neuron was calculated as

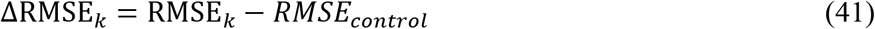

Two types of correlations are studied:

1. For a network (ANVN_R) with hidden neurons (*n* = 1: *N*), the Pearson correlation coefficient between ΔRMSE_*j*_ and *E_j_*, termed the Energy Error Correlation Coefficient (EECC), is calculated for each number of hidden neuron (n), where *E_j_* denotes the energy consumed by jth neuron, j ranging from 1 to N. The fig.16 shows the EECC between ΔRMSE and *E_j_* as the number of neurons is increased. For this network, the vascular network can return the excess energy to the reservoir. The network is simulated for different initial weights that connect the source node to the root node and reservoir. The results of all the simulations are averaged to get the plot shown in fig.1. The EECC was observed to be higher for a smaller number of hidden neurons, and as the number of hidden neurons increased, the EECC between the energy consumed by a neuron and its contribution to the network decreased slightly. This means that in larger networks, the neurons are not used as efficiently as in smaller networks. This seems to be the reason for a reduced efficiency observed for larger networks (fig 16.b) even though the maximum accuracy increased with the number of hidden neurons (fig 16.a).
2. For a network (ANVN) with a fixed number of hidden neurons N and variable input energy, where the network is unable to reject excess energy, the EECC between ΔRMSE_*j*_ and *E_j_* was calculated for each total input source energy *E_s_*, where the vector *E_j_* denotes the energy consumed by jth neuron, j ranging from 1 to N. The fig. 17 shows the correlation of ΔRMSE with *E_j_* as the input energy is increased. The number of leaf nodes was fixed at 512 hidden neurons. The network was simulated for different branching factors (K=2,3,4,6,8,16,32,64,256,512). The results of all the simulations are averaged to get the plot shown in fig.17 The EECC was higher for smaller energy, and as total input source energy increased, the EECC between the energy consumed by a neuron and its contribution to the network reduced significantly. This shows that when excess energy is available, the neurons are not efficiently utilized. In this network also, the lack of correlation of a neuron’s contribution to performance and its energy consumption might be the reason for a reduced efficiency when input energy is very high (fig 17.b).

**Figure 16:**
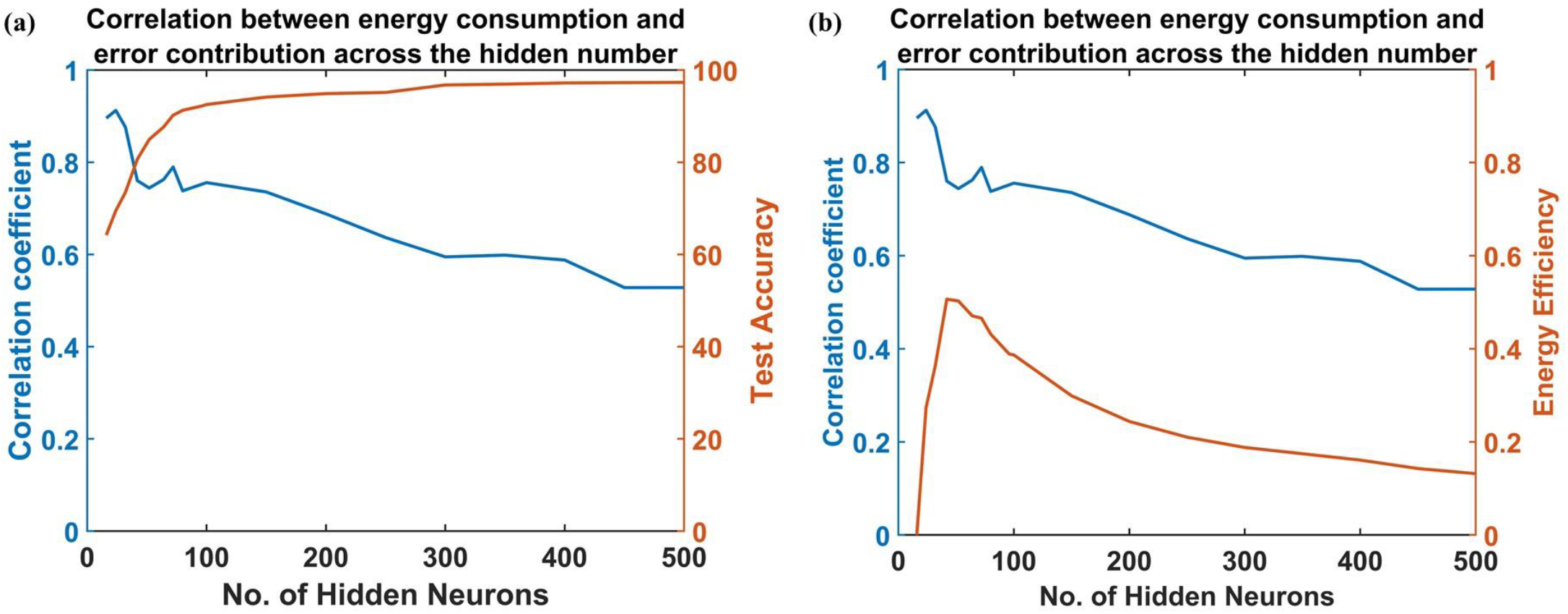
The variation of correlation coefficient with the number of hidden neurons. The network has the ability to return the excess energy to a reservoir. There is a high correlation at a low number of hidden neurons, and the correlation reduces as the number of neurons increases. (a) A comparison with change in accuracy. (b) A comparison with the change in energy efficiency.

**Figure 17:**
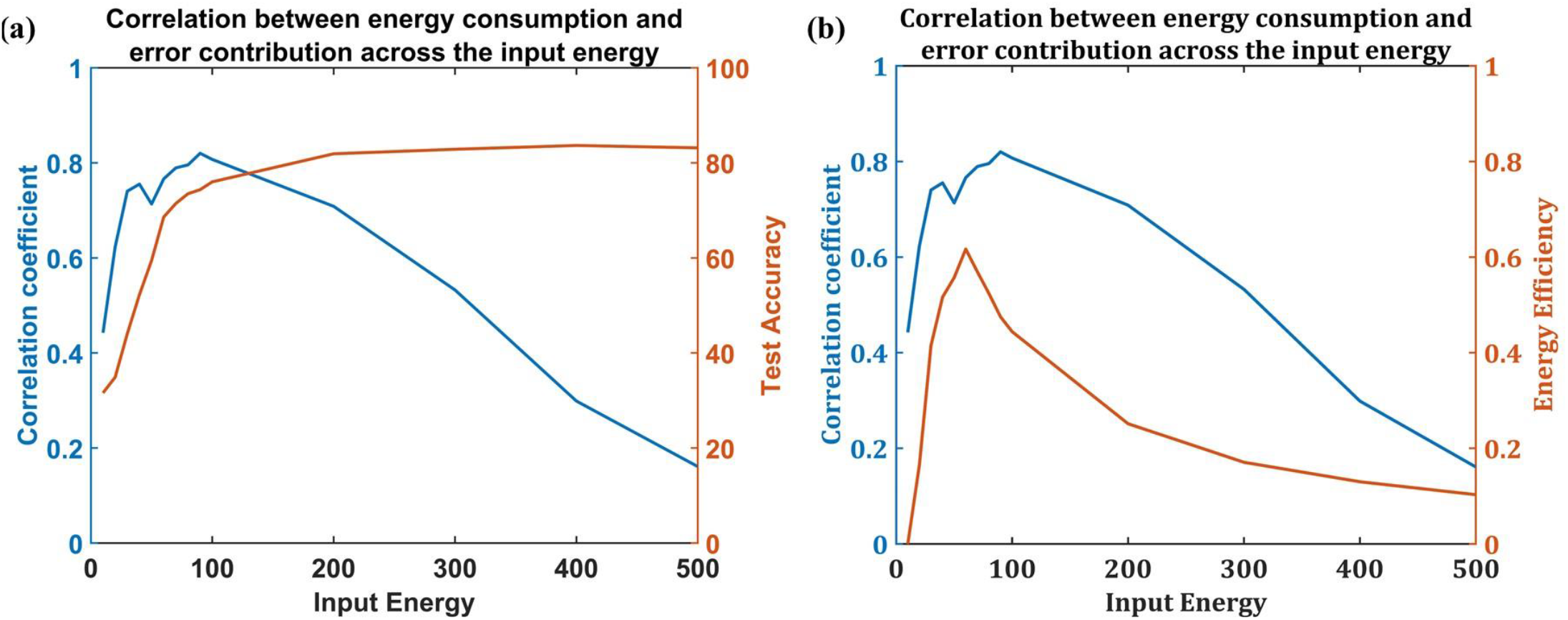
The variation of correlation coefficient with energy provided at the root node. The number of hidden neurons is fixed at N=512. There is no reservoir in the network, and the network cannot give back excess energy. There is a high correlation at a low number of hidden neurons, and the correlation reduces as the number of neurons increases. (a) A comparison with change in accuracy. (b) A comparison with change in energy efficiency.

## Discussion

One of the fascinating facts about the brain is that even though it only comprises 2% of the entire body mass, it consumes about 20% of the overall energy budget of the body. The evolution based on survival of the fittest placed a strict constraint on the available energy, which is reflected in the way the brain evolved (Aiello & Wheeler, 1995; Niven & Laughlin, 2008). Artificial neural networks inspired by biological neural networks have made significant advances in many fields and led to a renaissance of sorts in artificial intelligence (Chapter 6.3 in Hertz et al., 1991, Chapter 12 in Goodfellow et al., 2016). However, most artificial neural network models seem to focus on only information processing and ignore energy constraints. In this study, we tried to explore how energy dependence would affect the performance of an artificial neural network. Recent studies both experimental (Girouard, 2006; Hu et al., 2019; Iadecola, 2017; Owen & Sunram-Lea, 2011; Tarantini et al., 2017) and computational (Chander & Chakravarthy, 2012; Chhabria & Chakravarthy, 2016; Gandrakota et al., 2010; Kumar et al., 2018; Philips et al., 2016) have brought forth the importance of vascular and glial networks in the function of neural networks.

In this model, we simplified the complex energy delivery system of the biological neural network into an energy flow tree with weighted branches that are trainable. By comparing three training regimes where the vascular network was (i) untrained, (ii) sequentially trained after neural network, and (iii) simultaneously trained with the neural network, we show that simultaneous training of neural and the vascular network is the most energy-efficient. The training of vascular weights implies the rearrangement of the structure of the vasculature. The adaptation of the microvasculature following changes in the neural activity is well known and observed experimentally ^49,50^.

The ANVN_R showed that beyond a certain neural network size, the accuracy improvement was at the cost of energy efficiency. With the increase in the neural network size, the total energy consumption linearly increased while the increase in accuracy was negligible. Hence there exists a range of neural network sizes that can give an energy-efficient performance at a reasonable accuracy. In an environment where energy resources are limited, an ideal choice would be to settle for a network size where each neuron is maximally utilized, thereby permitting the neural network to be energy efficient. In a biological network, since the energy availability is limited, it makes sense that the network settles for a lower accuracy to achieve energy-efficient computing ^3,27,57,58^.

We explored the robustness of ANVN_R to the change in available initial energy. The network always converged to a stable settling point in the accuracy vs. per capita energy consumption space until a certain network size. Beyond that size, the trajectories of the neural network dynamics on the per capita energy consumption vs. accuracy space converged to a line of attractor, showing the dependence on the initial availability of the energy. This shows that the network size has to be small to ensure the robustness of the system. Providing an additional energy limitation constraint (*L*^1^ regularization) made the network more robust to the variation in initial energy available during training, and the stable settling point existed for a larger network size without compromising accuracy.

Retraining an ANVN_R with a different data set showed how transfer learning would manifest in the vascular network. The vascular weights at the level closest to the neurons (representing microvasculature) underwent maximum changes. Interpreting this result in biological terms, it appears that, in the vascular network, the retraining of the network using any new data set would result in a greater change at the level of the microvasculature than in larger vessels like the penetrating arterioles. This agrees with the microvascular plasticity observed in many in vivo models^59–63^.

Above all, the correlation between a neuron’ s energy consumption and its contribution to the network performance (quantified using EECC) was evident only when the network size was small. It was observed (Blue plot of fig 16 and 17) that when the number of hidden layers is small, there existed a positive correlation between the error contribution and energy consumption, meaning that the neurons that contribute the most towards the accuracy of the network consume more energy. However, surprisingly this correlation reduced with an increase in the number of neurons (fig. 16) as well as when the network receives higher energy (fig. 17). Even though the test accuracy for a network though achieved a higher value on increasing the number of hidden neurons (red plot in fig. 16.a) or input energy (red plot in fig. 17.a), this was at the cost of efficiency of the network (red plots in fig. 16.b and 17.b). This meant that at a larger network size, the neurons tend to be less efficient compared to the smaller network. The network ends up using all the available neurons instead of using just sufficient neurons, ensuring good performance but at the cost of very high energy consumption.

In this paper, we examine the relationship between energy availability and input/output performance in a MLP. It was earlier shown, using an electronic implementation of a Hopfield memory neural network, that improved retrieval performance is correlated with higher energy consumption in the form of increased dissipation through resistive elements ^64^. Similar results were also reported in the case of an oscillatory associative neural network model ^65^.

Taking a broader view, one must note that results that point to a link between energy consumption and informational performance are not limited to computational neuroscience models or even artificial neural network models. This link has been the central question of the field of the physics of computation and has a long history. Since irreversible processes in a thermodynamic system are associated with an irretrievable loss of energy as heat, more than half a century ago, Ralph Landauer asked if irreversible operations (e.g., addition: *x + y = z*) in a physical computing device are accompanied by dissipation of energy as heat. This profound question had led to the creation of the whole field of *reversible computation* ^66–68^. More recently, Karl Friston had proposed a free-energy theory of the brain that seeks to describe neuroenergetics and neural information processing in a unified thermodynamic framework ^69^. Theories of this type must be adequately extended and applied to a wide variety of computational neural architectures, both in neuroscience and in artificial intelligence, so as to simultaneously achieve an optimal informational and metabolic efficiency.

Our model presented the preliminary idea that the brain choosing energy-efficient performance over maximal performance is a characteristic of a robust neural network with limited energy availability. Extending this idea to a deep network would give an insight into how energy availability influences the feature extraction and learning exhibited by the neural networks. The correlation between the higher energy consumption levels and cognitive performance has become increasingly evident in recent years (Debatin, 2019, 2020; Geary, 2018, 2019, 2020; Lord et al., 2013; Owen & Sunram-Lea, 2011). Such a model would also help in understanding how energy availability impacts cognitive performance.

## Acknowledgments

We thank The Department of Biotechnology (DBT), Ministry of Science and Technology, Government of India for funding this project (BIO/17-18/303/DBTX/SRIN).

## Author Contributions

BSK did model designing, coding, analysis of the results, and manuscript preparation. NM did model designing and coding. SM did model designing and coding. VSC did model designing, analysis of the results, and manuscript preparation.

## Competing interests

The authors declare no competing interests.

## Notes

### Competing Interest Statement

The authors have declared no competing interest.

