## Supplementary Information for "Artificial Neurovascular Network (ANVN) to Study the Accuracy Vs. Efficiency trade-off in an Energy Dependent Neural Network"

### 1. The ANVN with reservoir (ANVN_R) results using EMNIST data set

The ANVN_R was simulated using the EMINST data also. The EMNIST is a dataset similar to MNIST, but with alphabets instead of numbers. We took 500 training data points and 200 test data points for all simulations, such that the data points were equally spanned in 10 predefined classes (Capital Letters A to J). The data was limited to 10 classes so as to use the same network of ANVN_R that we used for MNIST classification.

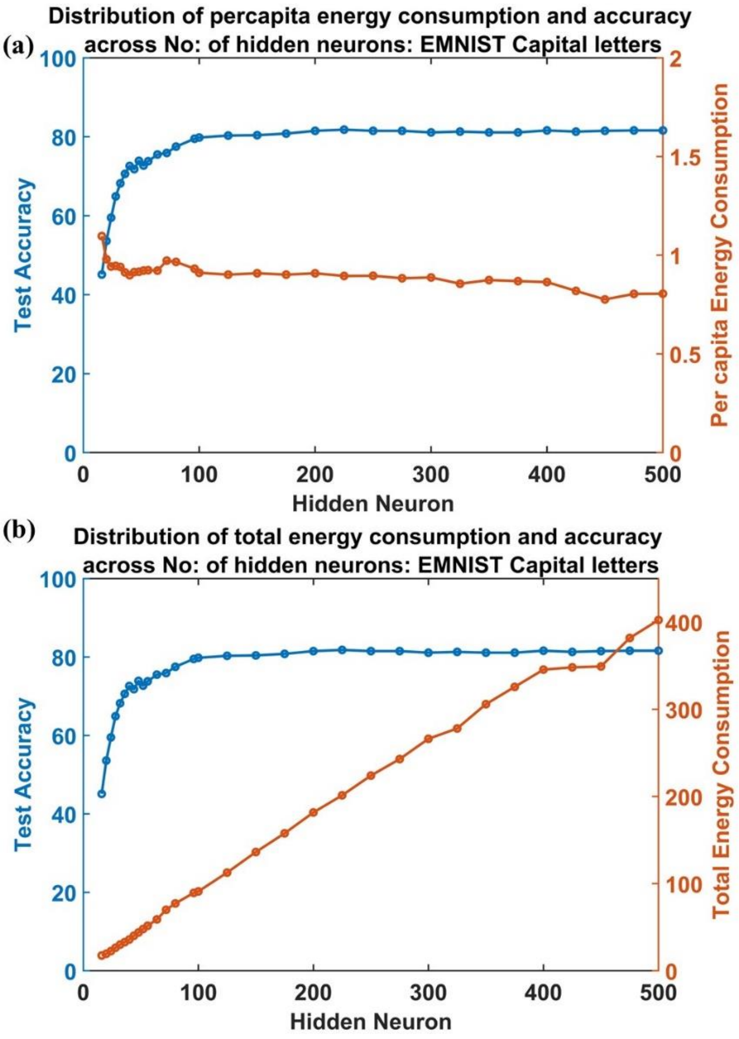


Figure S1: Study of variation in accuracy and energy consumption in ANVN_R with an increase in the number of hidden neurons (using EMNIST data set): (a) Test accuracy vs per capita energy consumption (b) Test accuracy vs total energy consumption

Figure S1.a,b shows the variation of percapita energy and total energy respectively as the number of neurons in the hidden layer increases. Similar to the results of MNIST, the EMNIST also gave a peak efficiency while the number of hidden neurons were around N=28 to N=36 (fig S2.a). Beyond that, the efficiency dropped and accuracy saturated. It was surprising that the robustness of the network to initial energy also was lost (fig S2.b) at almost similar network size (N~64) as obtained when run using MNIST

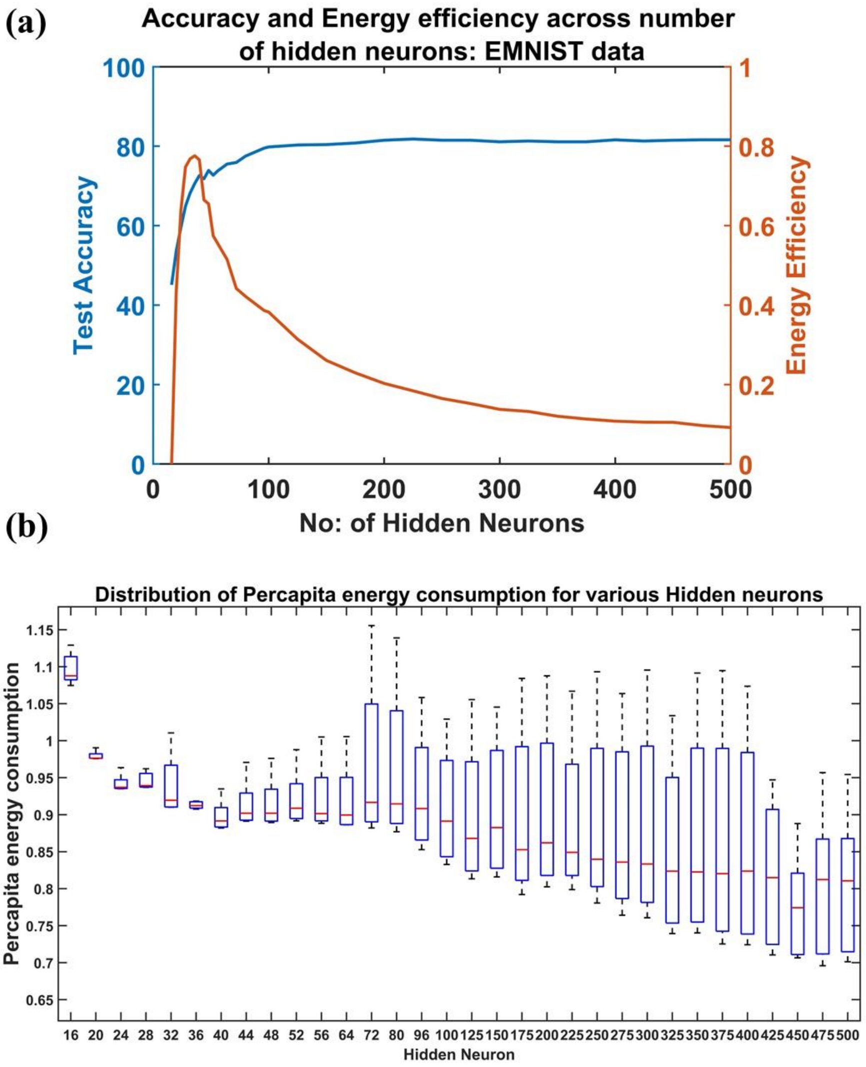


Figure S2: (a) The test accuracy and energy efficiency across the number of hidden layer neurons. (b) The box plot for visualizing the settling points of per capita energy consumption for each initial energy (varied from 0.2 units to 1 unit) given to single neurons given for each network size. The red mark shows the median value of the per capita energy consumed by the trained network and its maximum and the minimum values determine the height of the box (EMNIST data)

### 2. Transfer learning in ANVN trained under regime 3

The transfer learning was carried out in an ANVN network also and the results were similar to that of ANVN_R. As was observed in ANVN, the network learned faster when second data set was introduced (fig. S.3) and the difference in vascular weights was highest at the level closest to neurons (fig. S4 a,b). (Similar to the results obtained while using MNIST (fig.3 and fig 8.a), in the case of EMNIST also, the ANVN trained using regime 3 gave a lower accuracy than the ANVN with reservoir). The hidden layer had 512 neurons. Input energy given was 500 units. The branching factor was set to k=3 in order to simulate a network with 7 layers.

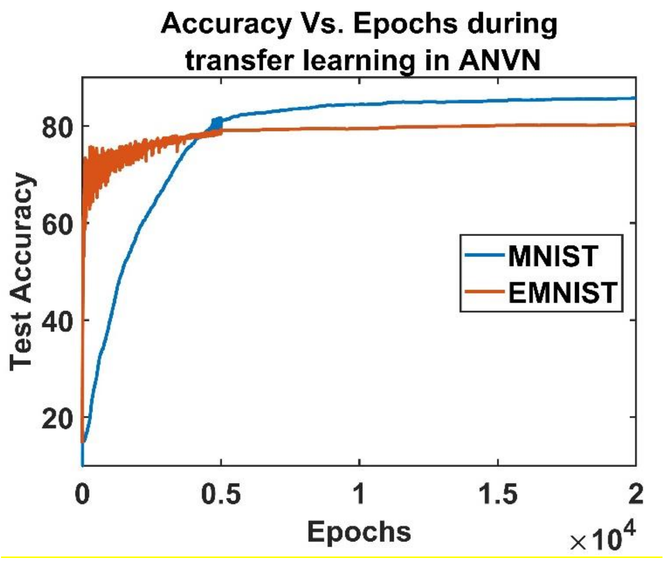


Figure S3: Training accuracy of the initially trained ANVN network (blue) takes more epochs to reach a high accuracy as compared to the network trained by transfer learning (red)

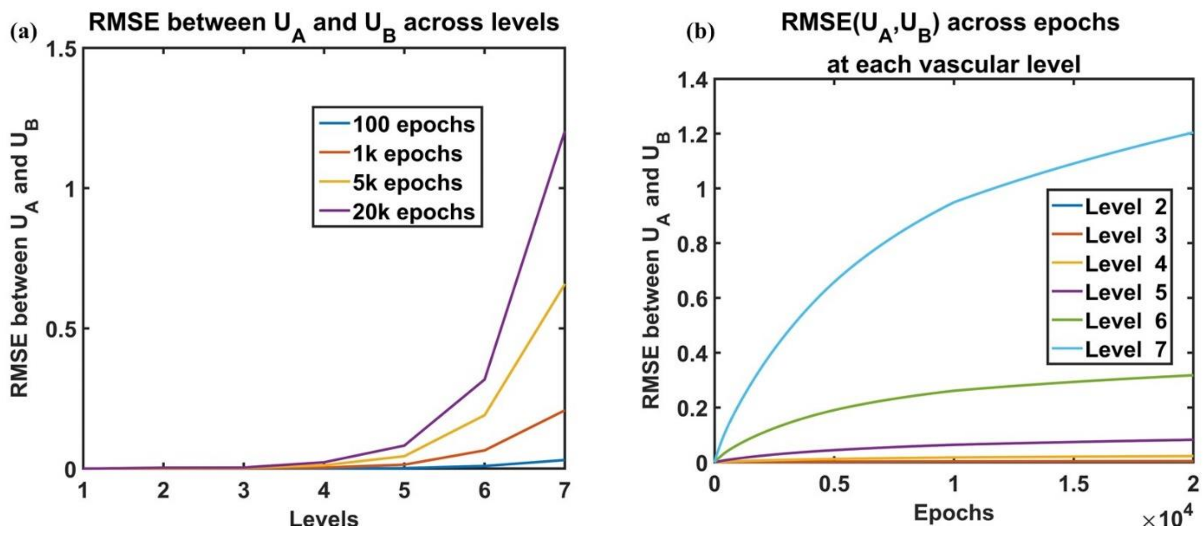


Figure S4: Variation of RMSE of the vascular weight at each level in the vascular tree (ANVN_R) (a) The RMSE between vascular weights at each level. The colors represent the RMSE at specified epochs. (b) The variation of RMSE across epochs for each level
